# General-purpose embeddings for long-read metagenomic sequences via *β*-VAE on multi-scale k-mer frequencies

**DOI:** 10.64898/2026.03.19.713080

**Authors:** Lauren M. Lui, Torben N. Nielsen

**Affiliations:** Environmental and Systems Biology Division, Lawrence Berkeley National Laboratory, Berkeley, CA 94720, USA; Jorgmundir, Tacoma, WA 98422, USA

**Author notes:** These authors contributed equally.

## Abstract

Long-read metagenomics routinely produces millions of assembled contigs, creating a need for methods that organize sequences into biologically meaningful groups across samples and environments. We present a general-purpose compositional embedding for metagenomic sequences based on a *β*-variational autoencoder (*β*-VAE) trained on multi-scale k-mer frequencies (1-mers through 6-mers; 2,772 features with centered log-ratio transformation). The embedding compresses each contig into a 384-dimensional vector that preserves local compositional similarity, enabling similarity search and graph-based clustering from sequence composition alone.

Through systematic comparison of fifteen models trained on up to 17.4 million contigs (525.5 Gbp) from brackish, terrestrial, and reference genome sources, we find that a small set of curated prokaryotic reference genomes (656,000 contigs) outperforms ten-fold larger domain-specific training sets, and that neither reconstruction loss nor Spearman correlation reliably predicts downstream clustering quality. On nearest-neighbor graphs, flow-based clustering (MCL) markedly outperforms modularity-based methods (Leiden), yielding 12,123 clusters from 154,041 contigs (≥ 100 kbp) with 99.2% phylum-level purity confirmed by independent marker gene phylogenetics. Multi-method taxonomic annotation achieves 87% coverage and reveals that 16.4% of contigs are eukaryotic—the single largest component invisible to standard prokaryotic annotation tools. The embedding provides a sample-independent coordinate system for organizing metagenomic sequence space at scale.

## Introduction

The advent of long-read sequencing technologies from Pacific Biosciences and Oxford Nanopore Technologies has transformed our ability to characterize microbial communities in complex environmental samples^1,2^. Short-read metagenomics produces highly fragmented assemblies in which individual genomes are scattered across many small contigs; long-read sequencing, by contrast, routinely generates contigs spanning tens to hundreds of kbp that can represent near-complete genomes, along with plasmids, viruses, and other mobile genetic elements^3,4^. This advance has opened new opportunities for understanding microbial ecology, metabolism, and evolution *in situ*, without the need for culturing. Yet because microbial communities harbor enormous diversity, individual long-read metagenomes still yield hundreds of thousands of assembled contigs, and environmental studies increasingly combine tens to hundreds of metagenomes spanning spatial or temporal gradients^5^. The combined assemblies can reach millions of contigs, and a critical challenge emerges: how to organize these sequences into biologically meaningful groups that reflect their genomic identities, both within and across samples.

Existing computational approaches to this problem focus on reconstructing individual genomes by grouping co-assembled fragments, typically combining compositional features with co-abundance information across multiple samples^6-8^. Long-read assemblies, however, increasingly produce contigs that span near-complete or complete genomes directly, reducing the need for computational genome reconstruction. More fundamentally, the goal of reconstructing a discrete, clonal genome may itself be ill-posed: in communities with high rates of recombination and gene flow, microbial populations can comprise clouds of related genotypes rather than distinct strains^9^. A different organizational challenge then arises: grouping compositionally similar sequences across samples and environments to reveal higher-level ecological and functional patterns—an approach that accommodates genomic variation within populations rather than forcing it into a consensus. Contigs from geographically distant sites may derive from organisms that are genomically close—potentially distinct species within the same lineage—and clustering by compositional similarity can provide an integrated view of community structure that transcends individual samples. This goal is distinct from metagenomic binning, which reconstructs individual genomes by combining composition with co-abundance across samples; we do not compare against binning tools, as they address a fundamentally different problem requiring different input data.

This cross-sample clustering requires a method for quantifying compositional similarity. Alignment-based methods identify sequences by homology to reference databases, but the majority of environmental microorganisms lack cultured representatives, and their sequences cannot be reliably classified through alignment alone^5^. Alignment-free approaches provide a scalable alternative, characterizing sequences by their composition alone without requiring homology to known references^10^. Genomic sequences carry species-specific compositional signatures, arising from factors such as codon usage bias, DNA structural constraints, and mutational biases^11,12^. Oligonucleotide frequencies provide a practical representation for encoding these signatures: by counting the occurrence of short subsequences (k-mers), each sequence can be represented as a fixed-length vector independent of sequence length or alignment. Tetranucleotide (4-mer) frequencies have become the standard choice for metagenomic applications^6-8,11^, offering 136 canonical features (unique k-mers after collapsing reverse complements) that balance discriminatory power with computational tractability. However, tetranucleotide frequencies capture only a limited view of sequence composition. They may not adequately distinguish closely related taxa with similar 4-mer profiles, nor do they capture the multi-scale structure inherent in genomic sequences—from dinucleotide biases reflecting DNA structural properties to hexanucleotide patterns shaped by restriction-modification systems. A representation that integrates compositional information across multiple scales could provide finer discrimination and support more flexible analyses.

Deep learning has emerged as a powerful paradigm for learning complex representations from high-dimensional biological data, with applications across genomics^13,14^. Variational autoencoders (VAEs)^15^, a class of generative models that learn compressed latent representations of data, combine dimensionality reduction with a regularized latent space that promotes smooth, continuous representations^16^. These properties make VAEs particularly suited to metagenomic sequence analysis: they require no taxonomic labels—essential when the majority of sequences lack reference representatives—and the regularized geometry of the latent space promotes meaningful distances between embeddings, enabling downstream tasks such as nearest-neighbor search and graph-based clustering directly in the learned representation.

Here, we develop a general-purpose embedding for long-read metagenomic sequences—one that is trained once and applied to any contig regardless of environment, organism, or domain of life—preserving local compositional similarity and enabling similarity search and graph-based clustering from a single representation. Because the embedding is derived entirely from sequence composition, it applies to any individual contig without requiring multi-sample experimental designs. We train *β*-VAE models^17^ on multi-scale k-mer frequencies (1-mers to 6-mers, 2,772 features with centered log-ratio (CLR) transformation^18,19^) from up to 17.4 million contigs (525.5 Gbp). Training data span brackish metagenomes from the San Francisco Estuary (SFE)^3^ and the Baltic Sea (SE; this study), soil, sediment, and water metagenomes from the Microflora Danica project^20^, and curated prokaryotic reference genomes from NCBI RefSeq.

Through systematic cross-comparison of fifteen models, we establish that a small, curated set of prokaryotic reference genome contigs can outperform much larger domain-specific training sets, and that neither reconstruction loss nor Spearman correlation reliably predicts downstream clustering quality—a finding underscored by the VAE’s superiority over PCA on taxonomic coherence despite PCA’s higher distance fidelity. On nearest-neighbor graphs where hub nodes often arise, flow-based clustering (MCL^21,22^) markedly outperforms modularity-based methods (Leiden^23^), yielding compositionally coherent clusters from 154,041 contigs (≥ 100 kbp). Independent taxonomic validation confirms >99% phylum-level cluster purity and reveals that a substantial fraction of contigs are eukaryotic—the single largest component invisible to standard prokaryotic annotation tools.

## Methods

### Datasets

We sought to organize and compare assembled contigs across 32 long-read metagenomes—16 from the San Francisco Estuary (SFE) and 16 from the Baltic Sea (SE)—to identify both shared and environment-specific genomic signatures at multiple scales: between individual samples, between the two environments, and across the combined dataset. To this end, we analyzed contigs from four sources spanning brackish aquatic, terrestrial, and reference genome origins.

#### San Francisco Estuary (SFE)

16 metagenomes from 8 sites, sampled in summer 2022 and winter 2023. Water samples were obtained from the USGS. DNA was extracted and sequenced on Oxford Nanopore PromethION (two flow cells per sample). Reads were assembled with Flye v2.9, yielding 2,467,377 contigs (59.9 Gbp) at ≥ 1 kbp. See Lui and Nielsen^1^ for details.

#### Baltic Sea (SE)

16 metagenomes from a summer 2023 transect across the Baltic Sea, in collaboration with Stefan Bertilsson (Swedish University of Agricultural Sciences). DNA was sequenced on Oxford Nanopore PromethION (two flow cells per sample) and assembled with Flye v2.9, yielding 4,226,452 contigs (70.7 Gbp) at ≥ 1 kbp.

#### Microflora Danica (FD)

154 samples (125 soil, 28 sediment, 1 water) from across Denmark, sequenced with Oxford Nanopore long reads and assembled as part of the Microflora Danica project. Contigs were prefiltered at 3 kbp before submission to the European Nucleotide Archive. The dataset comprised 9,817,648 contigs (295.7 Gbp) at ≥ 3 kbp. See Sereika *et al*.^2^ for details.

#### NCBI RefSeq Representative Genomes (NCBI)

A curated, dereplicated collection of ~20,000 representative genomes from NCBI RefSeq^3^, spanning bacterial and archaeal diversity. Downloaded December 7, 2025. The download comprised 1,117,682 contigs (99.3 Gbp); 903,568 (99.2 Gbp) at ≥ 1 kbp.

The focal datasets for downstream analysis are SFE and SE (32 brackish metagenomes, 6.7 million contigs). FD and NCBI were included in the training data exploration to test whether broader compositional diversity improves embedding quality; the rationale is described in Training Procedure below.

### K-mer Frequency Calculation and Preprocessing

Using a custom Python script, we extracted canonical k-mer frequencies for k = 1 through 6 from each contig in each of the four source datasets (SFE, SE, FD, NCBI) independently, producing one k-mer frequency matrix per dataset. This yielded 2,772 features per sequence: 2,080 hexamer (6-mer) frequencies, 512 pentamer (5-mer) frequencies, 136 tetramer (4-mer) frequencies, 32 trimer (3-mer) frequencies, 10 dimer (2-mer) frequencies, and 2 monomer (1-mer) frequencies. Frequencies were calculated by sliding a window along the forward strand and mapping each k-mer to its canonical form (the lexicographically smaller of the k-mer and its reverse complement), then normalizing by the total count within each k-mer size group, yielding compositional data that sum to one per group. We then pooled the per-dataset matrices and filtered by minimum contig length to create training subsets as needed (e.g., SFE+SE at ≥ 5 kbp for the SFE_SE_5 model, all four sources at ≥ 3 kbp for ALL_3; see Training Procedure). The full combination of all four sources comprised 17,415,045 contigs (≥ 1 kbp).

We chose to use multi-scale k-mer frequencies (1-mers to 6-mers) rather than a single k-mer size for several reasons. First, different k-mer sizes capture different aspects of sequence composition. Shorter k-mers (1-3) primarily reflect nucleotide composition and basic dinucleotide biases, mid-length k-mers (4-5) capture codon usage and local sequence context, and longer k-mers (6) provide more specific oligonucleotide signatures that can distinguish closely related taxa^4,5^. Second, the combination of multiple k-mer scales has been shown to improve taxonomic classification accuracy compared to single k-mer approaches^6,7^. Third, because 6-mers comprise the majority of input features (2,080 of 2,772), they are the most difficult to compress through the 384-dimensional bottleneck and dominate the reconstruction error (see Results), while shorter k-mers are more easily reconstructed and provide complementary signal. We did not include 7-mers despite their potential for finer-grained discrimination because: (1) they would nearly quadruple the input dimensionality (from 2,772 to 10,964 features), and (2) they are primarily useful for strain-level differentiation^8^ which was beyond our goal of species-level clustering.

K-mer frequencies represent compositional data, as they are constrained to sum to one^9^. Standard multivariate statistical methods assume data reside in Euclidean space and can lead to spurious correlations when applied to compositional data^10^. Because k-mer frequencies are normalized independently within each size group (each group sums to one), the k-mer frequency vector comprises six separate compositions. We therefore applied the centered log-ratio (CLR) transformation^11^ independently to each group, which removes the unit-sum constraint, mapping proportional data to Euclidean space:

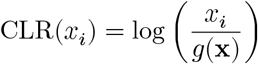

where g(x) is the geometric mean of all components within the group. To handle zero-count k-mers, we applied Jeffreys prior pseudocounts before transformation: for a group with *n* features, a pseudocount of 0.5/*n* was added to each frequency. This pseudocount is inspired by the Jeffreys prior Dir(0.5, …, 0.5)^12^, a standard uninformative prior for multinomial data, here applied to frequencies rather than counts. For hexamers (*n* = 2,080), this yields a pseudocount of 2.4 × 10^−4^, producing a gap of approximately 1.1 log units between zero-count and typical non-zero features. By contrast, a fixed pseudocount of 10^−6^ creates a gap of approximately 6 log units, causing the CLR geometric mean to be dominated by absent k-mers and obscuring frequency differences—a particular problem for shorter contigs where many 6-mers have zero counts. The CLR transformation has several advantages for our application: it removes the unit-sum constraint, making Euclidean distances meaningful in the transformed space; it reduces the dynamic range of the data; and it has been successfully applied in microbiome compositional data analysis^13,14^.

### Variational Autoencoder Architecture

We employed a *β*-Variational Autoencoder (*β*-VAE)^15,16^ to learn a lower-dimensional embedding of the CLR-transformed k-mer frequency vectors. The VAE framework^17^ learns both an encoder q_*φ*_(z|x) that maps input data x to a latent representation z, and a decoder p_*θ*_(x|z) that reconstructs the input from the latent code. The model is trained to maximize a modified evidence lower bound (*β*-VAE objective):

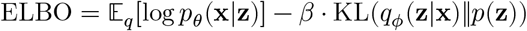

where the first term encourages accurate reconstruction, the second term regularizes the latent space to match a prior distribution p(z) = N(0, I), and *β* controls the trade-off between the two objectives. Our implementation minimizes the negative of this objective, averaged over each mini-batch of N samples:

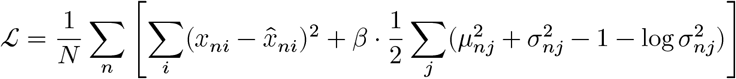

where i indexes the 2,772 CLR-transformed input features, j indexes the 384 latent dimensions, and *β* = 0.05. The reconstruction term is the sum of squared errors (SSE) across all 2,772 features—not the per-feature mean—while the KL term sums over 384 latent dimensions and is down-weighted by *β* = 0.05; this formulation strongly favors reconstruction. Tables report per-feature MSE (SSE/2,772) for readability; reproducing our results requires matching this SSE-based loss formulation.

Two properties of the VAE are central to our application. First, the KL regularization term encourages the encoder to produce overlapping posterior distributions, enforcing continuity in the latent space—precisely the property needed for meaningful nearest-neighbor retrieval. Second, stochastic sampling during training exposes the decoder to a different perturbation of each latent encoding at every epoch, effectively multiplying the variety of training inputs. Together with the KL penalty, this prevents the model from memorizing individual input–output pairs, instead forcing it to learn the underlying distribution of the data. The reparameterization trick^17^ makes this sampling step differentiable, enabling gradient-based optimization of both properties. Training for 1,000 epochs reinforces these effects: each of the training sequences is encoded and decoded with different random perturbations a thousand times, progressively smoothing the mapping and allowing even a relatively small number of training points to populate the latent space with coherent regions.

Our architecture follows a shallow, symmetric design. We chose progressive dimensionality reduction through the encoder (2,772 → 1,024 → 512 → 384) rather than a single abrupt bottleneck, and focused optimization effort on the latent dimensionality and *β*, which most directly affect embedding quality (see below). The encoder consisted of two fully-connected hidden layers with 1,024 and 512 units, respectively, each followed by batch normalization^18^ (to stabilize training across large batches) and Leaky ReLU activation (*α* = 0.2). The encoder produced two 384-dimensional vectors representing the mean (*μ*) and log-variance (log *σ*^2^) of the latent distribution. The log-variance was clipped to the range [–20, 2] to prevent numerical instability during the reparameterization step. We used the reparameterization trick^17^ to sample from the latent distribution during training: z = *μ* + *σ* ⊙ *ε*, where *ε* ~ N(0, I).

The decoder mirrored the encoder architecture, with 512 and 1,024 unit hidden layers (again with batch normalization and Leaky ReLU), followed by a final 2,772-unit linear output layer to reconstruct the CLR-transformed k-mer frequencies. The total model contained approximately 7.3 million trainable parameters.

### Choice of Latent Dimensionality and *β*

The latent dimensionality and the *β* hyperparameter critically affect VAE performance. A small latent dimension may be insufficient to capture the complexity of metagenomic sequence space, while an excessively large latent space may fail to learn useful structure and can be computationally expensive for downstream clustering. Similarly, *β* controls the balance between reconstruction fidelity and latent space regularization: with *β* = 1, the KL regularization is relatively stronger and can restrict the encoder’s expressiveness^19,20^, while very low *β* (approaching 0) creates an irregular autoencoder without the beneficial structure imposed by the KL term. Setting *β* < 1 allows the encoder to use the latent space more expressively for local distance preservation while retaining the regularization that prevents degenerate solutions.

We performed a systematic hyperparameter search over latent dimension ∈ {256, 384} and *β* ∈ {0.03, 0.05, 0.1, 0.2}, training each configuration for 500 epochs on the ALL_5 dataset (13,439,298 contigs from all four sources at the ≥ 5 kbp threshold). We evaluated models based on four criteria: (1) reconstruction mean squared error (MSE) in CLR space, (2) KL divergence magnitude (as a proxy for posterior collapse—very low KL indicates an uninformative latent space), (3) training stability (convergence without oscillation), and (4) local distance preservation measured by Spearman correlation between latent-space distances and k-mer MSE (see Embedding Quality Evaluation).

The best configuration we identified was latent dimension = 384 and *β* = 0.05. This model consistently achieved the lowest reconstruction MSE across *β* values, maintained healthy KL divergence well above levels indicative of posterior collapse, and exhibited stable training convergence. In comparison, *β* = 0.03 showed less stable training with higher-variance validation metrics, while *β* = 0.1 and 0.2 produced worse reconstruction without meaningful improvements in latent space structure. The 256-dimensional latent space, even at the best *β*, showed higher 6-mer reconstruction error compared to the 384-dimensional space (see Results), suggesting that the additional dimensions were needed to adequately represent hexamer diversity. This configuration was subsequently validated across all five minimum contig length thresholds (see Results).

The relatively low *β* value (0.05 compared to the standard *β* = 1) reflects the specific requirements of our application. Our primary goal was to preserve local distances in the embedding space to enable accurate nearest-neighbor retrieval and clustering. The KL term in VAEs encourages the latent distribution to match the prior N(0, I), which can smooth over local structure in favor of global regularization. By reducing *β*, we allowed the encoder to use the latent space more expressively to preserve local relationships while still maintaining sufficient regularization to avoid overfitting (as evidenced by similar training and validation losses). This design choice is consistent with previous work showing that the optimal *β* depends on the downstream task, and that values below 1 can improve representation quality when local distance preservation matters more than global regularization^21,22^.

### Training Procedure

Short contigs yield noisier k-mer frequency estimates because fewer k-mer positions are observed relative to the number of possible k-mers. For example, a 1 kbp contig provides approximately 995 canonical hexamer observations across 2,080 possible canonical 6-mers, leaving many k-mers with zero counts.

Our downstream clustering analysis focuses on the brackish metagenomes (SFE and SE). However, we hypothesized that including additional training data from diverse environments would improve embedding quality by exposing the model to a broader range of compositional patterns. To test the effects of both minimum contig length and training data composition on embedding quality, we trained fifteen VAE models, each on a distinct dataset defined by its source combination and minimum contig length threshold (Table 1). Because filtering by length changes the size, composition, and length distribution of the training data, each threshold produces an effectively different dataset (e.g., SFE_SE_2 contains 6.5 million contigs while SFE_SE_5 contains 4.8 million with a median length of ~8 kbp).

**Table 1:** Dataset sizes across minimum contig length thresholds.

| Min length | SFE | SE | FD | NCBI | ALL total | Brackish total |
| --- | --- | --- | --- | --- | --- | --- |
| 1 kbp | 2,467,377 | 4,226,452 | 9,817,648 | 903,568 | 17,415,045 | 6,693,829 |
| 2 kbp | 2,390,156 | 4,064,984 | 9,817,648 | 782,906 | 17,055,694 | 6,455,140 |
| 3 kbp | 2,319,168 | 3,632,008 | 9,817,648 | 726,950 | 16,495,774 | 5,951,176 |
| 4 kbp | 2,112,286 | 3,196,949 | 8,826,953 | 686,768 | 14,822,956 | 5,309,235 |
| 5 kbp | 1,941,549 | 2,835,221 | 8,006,888 | 655,640 | 13,439,298 | 4,776,770 |

The first ten models comprised two source combinations at five length thresholds (1, 2, 3, 4, and 5 kbp). ALL models included all four datasets (SFE, SE, FD, NCBI) to maximize compositional diversity (13.4– 17.4 million contigs), with FD broadening the compositional space beyond brackish environments and NCBI reference genomes providing taxonomic signposts—sequences of known origin that anchor the embedding space. SFE_SE models included only the focal brackish metagenomes to test whether domain-specific training produces better embeddings for the target analysis domain (4.8–6.7 million contigs).

Based on the initial cross-comparison results (see Results), we trained five additional models to test specific hypotheses about how training data composition, source diversity, and length distribution affect downstream clustering performance on long contigs (≥ 100 kbp). These were: NCBI_5 (NCBI RefSeq only, ≥ 5 kbp; 655,640 contigs from ~20,000 representative genomes), NCBI_100 (NCBI RefSeq only, ≥ 100 kbp; 175,213 contigs), SFE_SE_100 (brackish only, ≥ 100 kbp; 154,041 contigs), ALL_100 (all four sources, ≥ 100 kbp; ~845,000 contigs), and SFE_SE_NCBI_5 (brackish + NCBI, ≥ 5 kbp; ~5.4 million contigs). All fifteen models used identical architecture and hyperparameters.

Note that the Microflora Danica (FD) contigs were pre-filtered at 3 kbp before submission to ENA, so the 1 and 2 kbp ALL thresholds gain additional shorter contigs only from SFE, SE, and NCBI sources.

For each model, we concatenated the source k-mer frequency matrices, randomly shuffled the combined dataset (seed 42), and split into training (90%) and validation (10%) sets.

Each model was trained using the Adam optimizer^23^ with an initial learning rate of 10^−4^, *β*_1_ = 0.9, *β*_2_ = 0.999, and batch size of 1,024. We employed ReduceLROnPlateau scheduling, halving the learning rate when validation loss failed to improve for 20 consecutive epochs, down to a minimum of 10^−6^. To prevent early-training posterior collapse, we used KL warmup^19^: *β* was linearly annealed from 0 to its target value (0.05) over the first 5 epochs, allowing the decoder to learn meaningful reconstructions before the full KL penalty was applied. Each model was trained for 1,000 epochs, saving checkpoints whenever validation loss improved.

Training was performed using Keras 3.12 with the JAX 0.8.1 backend (Python 3.12) on a system equipped with a 32-core AMD Ryzen Threadripper PRO 7975WX CPU, an NVIDIA RTX 6000 Ada Generation GPU, and 512 GB of DDR5 RAM. Training runs of 1,000 epochs required 7–11 hours depending on dataset size (139–180 GB peak memory for the k-mer matrices alone). Batch normalization running statistics were updated during training but frozen during validation and inference.

The best model for each configuration (selected by minimum validation loss) was used to generate embeddings by encoding contigs through the trained encoder, using the deterministic latent mean (*μ*) rather than the stochastic sample (z) to ensure reproducible embeddings.

While we used dedicated hardware, none of the training runs described here would exceed $20 at current cloud GPU pricing, making this approach accessible to groups without local GPU infrastructure.

### Embedding Quality Evaluation

Our goal is not to identify an optimal VAE architecture, but to produce embeddings that are useful for organizing metagenomic contigs. We therefore evaluate embedding quality by the fidelity of local distance relationships—whether sequences similar in k-mer composition are close in latent space—rather than by reconstruction loss alone, since downstream clustering and retrieval depend on distance-based comparisons. We validated this property using the following protocol:

1. Loaded 50,000 contigs from the start of the held-out validation set (the last 10% of each training file, the same split used for monitoring training loss and learning rate scheduling). The pool size was standardized to 50,000 across all evaluations because Spearman correlation is sensitive to pool size—larger pools provide more neighbor candidates at diverse distances, inflating the correlation. Standardizing ensures fair comparison across datasets of different sizes.
2. Encoded all contigs using the trained encoder (deterministic z_mean)
3. Selected 100 random queries from the 50,000 samples
4. For each query, found the 50 nearest neighbors in latent space using Euclidean distance
5. For each query-neighbor pair, computed the ground-truth k-mer dissimilarity as the MSE between their CLR-transformed k-mer frequency vectors
6. Calculated Spearman rank correlation between latent Euclidean distances and k-mer MSE across all query-neighbor pairs
7. Generated a random baseline by computing k-mer MSE for random pairs

We used Spearman rank correlation (*ρ*) as the primary evaluation metric because it measures the rank-ordering fidelity of the embedding—whether closer points in latent space are genuinely more similar in k-mer composition—independent of the scale or functional form of the distance relationship. This is the property that matters for nearest-neighbor retrieval and graph-based clustering. The correlation is computed over all 5,000 query-neighbor pairs pooled together (100 queries × 50 neighbors). Pooling across queries from regions of varying density may inflate absolute Spearman values compared to per-query estimates, but because all models are evaluated using the identical protocol (same queries, same pooling), relative comparisons across models are unaffected. We quantified uncertainty by bootstrap resampling over queries (10,000 resamples), yielding 95% percentile confidence intervals of approximately $±$0.06, reflecting the 100-query sampling unit.

We chose Euclidean distance over cosine distance for latent-space comparisons based on a direct comparison showing that cosine distance suffers from concentration of distances in this high-dimensional space^24^ (see Results). All subsequent analyses used Euclidean distance.

This protocol was applied across all fifteen models in a full cross-comparison design. To enable fair comparison across models trained on different data distributions, each model was evaluated not only on its own held-out data but on the held-out sets from all five models within its family (ALL or SFE_SE), producing 5$×$5 Spearman correlation matrices. In cross-model evaluations (off-diagonal entries), the held-out data was never seen during training of the evaluated model, providing a genuine test of generalization. Additionally, ALL models were evaluated on all SFE_SE held-out sets and vice versa, enabling cross-domain comparison. The five experimental models were evaluated alongside the best and worst ALL models (ALL_3 and ALL_5) and the best brackish model (SFE_SE_1) on brackish held-out data (SFE_SE_5 and SFE_SE_100) and NCBI held-out data (NCBI_5), producing an 8-model × 3-dataset comparison matrix (see Results). We use “test data” in Results tables as shorthand for these held-out sets. On-diagonal entries (a model evaluated on its own data) use the same validation split that guided model selection; off-diagonal entries are a genuine test of generalization.

### Baseline Comparisons

To contextualize the VAE embedding, we compared it against two linear baselines: PCA at 384 components (matching the VAE latent dimensionality) and raw CLR-transformed features at the full 2,772 dimensions (an upper bound on distance fidelity, as the Spearman evaluation metric measures rank concordance with CLR-space MSE). PCA was fitted on the same NCBI_5 training split (90%, 500,000 randomly sampled sequences) and applied to the 154,040 contigs at ≥ 100 kbp using scikit-learn^44^.

Both baselines were processed through the identical downstream pipeline: 50-nearest-neighbor search using Euclidean distance, graph construction with distance threshold calibrated to match connectivity (nn1 percentile matching to the VAE graph’s 86.8% connectivity), in-degree capping at 100, and MCL clustering at I = 3.0. Taxonomic evaluation used V-measure, adjusted Rand index (ARI), normalized mutual information (NMI), and pairwise precision/recall/F1 computed from GTDB-Tk direct classifications (26,841 classified contigs at ≥ 100 kbp), with completeness analysis measuring the fraction of each taxon’s members contained within a single cluster.

### Embedding Generation and Nearest-Neighbor Search

Based on the cross-comparison and downstream clustering results (see Results), we selected the NCBI-only model trained at the 5 kbp threshold (NCBI_5) for downstream clustering.

We generated embeddings for all 6,693,829 SFE and SE contigs at ≥ 1 kbp by passing their CLR-transformed k-mer frequency vectors through the trained encoder, using the deterministic latent mean (*μ*). The resulting 6,693,829 × 384 matrix (~10 GB in float32)—a 7.2-fold reduction in dimensionality from the 2,772-dimensional input—was loaded into a ChromaDB 1.3.5^25^ vector database configured with L2 (Euclidean) distance and HNSW (Hierarchical Navigable Small World) indexing^26^ (default parameters: M = 16, ef_construction = 100) for efficient approximate nearest-neighbor search. HNSW enables sub-linear query time, finding the k = 50 nearest neighbors among 6.7 million embeddings in milliseconds on a single CPU core.

For clustering analysis, we applied minimum contig length filters from 3 to 500 kbp to assess the relationship between sequence length and clustering quality. At each threshold, we queried ChromaDB for the k = 50 nearest neighbors of each sequence, restricting the search to sequences meeting the length threshold. The resulting neighbor lists, with associated Euclidean distances, formed the basis for graph construction.

### Graph Construction for Community Detection

We constructed nearest-neighbor graphs from the query results using three strategies to address the hub problem—the tendency for certain sequences in high-dimensional spaces to appear as nearest neighbors of disproportionately many other sequences^27,28^.

#### Symmetric k-NN graph

For each pair (i, j) where j appears in i’s k-nearest-neighbor list or i appears in j’s, we added an undirected edge. This maximizes coverage but allows hub nodes to create giant connected components through transitivity chains: if sequences A and B both point to a shared hub, the symmetric graph connects A and B through the hub even if they are compositionally dissimilar.

#### Mutual k-NN graph

Edges were retained only when both directions existed: j must be among i’s k-nearest neighbors and i among j’s. This eliminates hubs because a hub’s own nearest neighbors are typically closer to each other than to the many distant sequences pointing at the hub. However, mutual filtering severely reduces coverage by isolating the majority of sequences.

#### In-degree capped graph

Starting from the directed k-NN graph, we capped the in-degree of each node at 100, retaining only the 100 shortest-distance incoming edges when a node’s in-degree exceeded the cap, then symmetrized the result. The cap was set well above the 99th percentile of in-degree in dense regions, so it selectively removed only extreme hubs while preserving graph structure for the vast majority of sequences.

Edges were weighted using the distance-based function w(d) = 1/(d + 0.1), which provides a gentle similarity measure that decreases with increasing Euclidean distance. We also tested exponential weighting w(d) = exp(–d), but found it substantially worse for MCL clustering: exponential weighting over-suppresses medium-range edges (d = 3–5) that MCL’s flow simulation needs for cluster boundary definition (see Results).

We applied a Euclidean distance threshold to remove edges between sequences exceeding a maximum distance. The threshold was selected based on nearest-neighbor distance analysis: at each length filter, we computed the distance to the single nearest neighbor (nn1) for a random sample of 10,000 sequences. The optimal distance threshold was chosen to include the majority of genuine nearest-neighbor pairs while excluding noise (see Results for the nn1 sweep across length thresholds).

### Clustering Methods

We applied two community detection algorithms to the weighted nearest-neighbor graphs: the Markov Cluster Algorithm (MCL v22-282)^29,30^ as the primary clustering method, and the Leiden algorithm^31^ for comparison.

**MCL** simulates random walks on the graph through alternating expansion (matrix squaring) and inflation (entrywise power followed by column renormalization) operations^29^. The expansion step propagates flow across paths of length 2, while the inflation step amplifies strong connections and attenuates weak ones. After convergence, flow circulates within clusters and dies between clusters, naturally identifying community structure. The inflation parameter I controls cluster granularity: higher values produce smaller, tighter clusters. We tested I ∈ {1.4, 2.0, 3.0, 4.0, 5.0} to characterize the granularity trade-off.

**Leiden**^31^ optimizes a modularity-based quality function to partition the graph into communities with high internal edge density and low inter-community connectivity. We used the leidenalg v0.11.0 implementation (igraph 1.0.0) with the modularity quality function (RBConfigurationVertexPartition, the default). We tested resolution parameters spanning a 10-fold range (0.2, 0.3, 0.5, 0.7, 1.0, 1.3, 1.6, 2.0); Leiden produced nearly identical partitions across this range, indicating insensitivity to the resolution parameter on this graph (see Results for cluster counts). We report results at resolution 1.0.

MCL was chosen as the primary method because its flow-based approach naturally attenuates weak transitive connections. If sequence A connects to a hub which connects to unrelated sequence B, the indirect flow A → hub → B is progressively weakened by inflation at each iteration. By contrast, modularity optimization maximizes internal edge density relative to a null model; this approach can merge compositionally dissimilar nodes connected through shared hubs into a single community, making Leiden vulnerable to the transitivity chain problem on metagenome graphs where hub nodes arise.

### Cluster Quality Assessment

We assessed cluster quality using GC content span as the primary metric. For each cluster, we computed the range of GC content (maximum minus minimum) across all member sequences, expressed in percentage points (pp). GC content is a strong proxy for taxonomic coherence: sequences from the same species share characteristic GC content arising from mutation bias and selection^32^, so clusters with narrow GC spans are more likely to represent genuine biological groups. We report GC spans for the three largest clusters at each configuration as representative quality indicators, along with the median, 95th percentile, and maximum GC span across all non-singleton clusters.

We estimated the intrinsic dimensionality of the embedding manifold using the TWO-NN estimator^33^, which uses only the ratio of second-to-first nearest-neighbor distances to estimate the local dimensionality of a dataset without density assumptions. This metric characterizes the geometric complexity of the data at each length threshold, providing insight into the effective dimensionality of the manifold on which the embeddings reside despite the 384-dimensional ambient space.

To determine the optimal minimum contig length for clustering, we conducted a nearest-neighbor distance (nn1) sweep: for each minimum length from 3 to 500 kbp, we randomly sampled 10,000 sequences and computed the Euclidean distance to each sequence’s single nearest neighbor. The distribution of nn1 values reveals whether sequences at a given length threshold form dense neighborhoods (low nn1, amenable to clustering) or are isolated singletons (high nn1, indicating compositional isolation from any clustered population).

To provide a theoretical basis for the observed length threshold, we modeled k-mer frequency precision using multinomial sampling statistics. Under a multinomial model, the coefficient of variation of each canonical k-mer’s frequency estimate scales approximately as 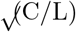, where C is the number of canonical k-mer types in the group and L is the contig length in base pairs (exact: 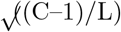; the approximation is tight for C ≫ 1, which holds for all groups except 1-mers). This predicts the minimum length at which k-mer frequencies become precise enough for reliable compositional comparison.

We visualized the global structure of the latent space using t-SNE^34^ (openTSNE 1.0.4, perplexity = 30, early exaggeration = 12) to examine the organization of embeddings with respect to GC content and dataset of origin.

### Taxonomic Validation

Based on the nearest-neighbor distance sweep (see Results), we selected 100 kbp as the minimum contig length for downstream clustering and validation—a threshold that is practical specifically because long-read metagenomics routinely produces contigs of this length. To assess whether the compositional clusters identified by MCL correspond to biologically coherent groups, we applied five independent annotation methods— NCBI signpost analysis, GTDB-Tk marker gene phylogenetics, MMseqs2 protein-level taxonomy, geNomad virus and plasmid detection, and Tiara eukaryotic domain classification—plus taxonomy propagation within MCL clusters, supplemented by coding density analysis for independent corroboration. All methods were applied to the full set of 154,041 contigs (≥ 100 kbp) from the SFE and SE metagenomes.

#### NCBI signpost analysis

In addition to the 154,041 brackish contigs, we embedded all 175,213 NCBI RefSeq representative genome contigs (≥ 100 kbp) through the trained NCBI_5 encoder and identified the nearest brackish contig to each reference sequence in the 384-dimensional latent space. Reference–brackish pairs within Euclidean distance d < 5.0 were retained as taxonomic signposts. We then propagated the NCBI taxonomy—retrieved programmatically from the NCBI Taxonomy database via Entrez—to brackish contigs sharing a cluster with at least one signpost. Cluster-level consensus was determined by majority vote: a taxonomic rank was assigned to the cluster only when ≥ 80% of signpost-linked members agreed.

#### GTDB-Tk marker gene phylogenetics

We ran GTDB-Tk v2.6.1^35^ on all 154,041 brackish contigs using the GTDB r226 reference database^36^. GTDB-Tk identifies 120 bacterial and 53 archaeal marker genes via hidden Markov models and places sequences into the GTDB reference tree using pplacer. We used the --skip_ani_screen flag to bypass the ANI-based screening step, which is designed for complete genomes rather than metagenomic contigs. This method is methodologically independent of k-mer composition: it relies on protein sequence homology of conserved marker genes rather than oligonucleotide frequencies. We compared GTDB-Tk and NCBI signpost assignments for cross-validated contigs to assess inter-method agreement, applying 73 explicit name mappings to reconcile NCBI and GTDB nomenclature differences (e.g., NCBI Betaproteobacteria → GTDB Gammaproteobacteria).

#### MMseqs2 protein-level taxonomy

We independently classified all contigs using MMseqs2 v18.8cc5c^37,38^ taxonomy against the NCBI non-redundant protein database (nr, downloaded February 2026)^39^. ORFs were predicted internally by MMseqs2, and each ORF was assigned taxonomy using the two-step best lowest common ancestor (2bLCA) protocol (`-s 4.0, --lca-mode 2`), which identifies the best hit, realigns its matched region against all candidate homologs, and computes the lowest common ancestor of all hits with E-values below that of the best hit. We aggregated ORF-level assignments to contig-level consensus: at each taxonomic rank, the contig was assigned the majority taxon only when ≥ 80% of classified ORFs agreed— the same 80% agreement threshold used for NCBI signpost consensus. This method is complementary to both GTDB-Tk (which relies on 120/53 conserved marker genes) and the k-mer embedding (which uses oligonucleotide composition): MMseqs2 queries all predicted proteins against a comprehensive reference database, providing broader taxonomic coverage at the cost of higher computational requirements.

#### Taxonomy propagation within MCL clusters

We propagated GTDB-Tk taxonomy from classified contigs to unclassified cluster-mates. For each cluster with ≥ 2 GTDB-Tk-classified members, we assigned taxonomy to unclassified members by majority vote, requiring ≥ 80% agreement at each rank. This propagation is justified only if clusters are taxonomically coherent; we assess this in Results.

#### geNomad virus and plasmid detection

We screened all contigs using geNomad v1.11.2^40^, which combines a neural network classifier operating on raw DNA sequence with a separate marker gene-based classifier to identify viral and plasmid sequences. geNomad is complementary to GTDB-Tk: it targets mobile genetic elements that lack the conserved marker genes used for prokaryotic phylogenetics.

#### Tiara eukaryotic domain classification

We ran Tiara v1.0.3^41^, a deep learning classifier trained to distinguish bacterial, archaeal, eukaryotic, organellar, and unknown sequences based on k-mer frequencies. Tiara fills a critical gap: GTDB-Tk classifies only prokaryotes, geNomad detects only viruses and plasmids, and NCBI signposts are limited to prokaryotic reference genomes.

#### Coding density analysis

We predicted protein-coding genes on all contigs using pyrodigal v3.6.3^42^ (a Python implementation of Prodigal^43^) in metagenomic mode. Coding density—the fraction of each contig covered by predicted coding sequences—provides independent corroboration of domain assignments: prokaryotes, eukaryotes, and giant viruses occupy distinct ranges of gene organization, enabling cross-referencing with Tiara and geNomad classifications.

#### Cluster purity assessment

For each MCL cluster with ≥ 2 taxonomically annotated members, we computed purity as the fraction of members sharing the majority taxon at each rank (domain through genus). A cluster was considered perfectly pure at a given rank if all annotated members agreed. We report both mean purity and the percentage of perfectly pure clusters across all annotated clusters.

## Results

### Systematic Optimization of VAE Hyperparameters

To develop an effective embedding for metagenomic sequences, we explored the VAE architecture through systematic hyperparameter evaluation using the ALL_5 dataset (13,439,298 contigs from all four sources at the ≥ 5 kbp threshold). We trained models with varying latent dimensions (256 and 384) and *β* values (0.03, 0.05, 0.1, 0.2), evaluating each on four key criteria: reconstruction accuracy (MSE in CLR-transformed k-mer space), KL divergence magnitude as an indicator of latent space expressiveness, training stability, and local distance preservation measured by Spearman correlation between latent Euclidean distances and k-mer dissimilarity.

The 384-dimensional latent space consistently outperformed the 256-dimensional space across all *β* values, reducing 6-mer reconstruction MSE by approximately 13% while maintaining similar KL divergence values— suggesting that the additional dimensions were necessary to adequately capture the diversity of hexamer patterns across the four data sources spanning brackish, non-brackish, and reference genome origins. Across *β* values, we observed a clear trade-off between reconstruction fidelity and latent space regularization: lower *β* values (0.03, 0.05) achieved better reconstruction with higher KL divergence, while higher *β* (0.1, 0.2) showed worse reconstruction with more regularized latent spaces. All configurations maintained KL divergence well above the near-zero levels indicative of posterior collapse. *β* = 0.03 showed less stable training with higher variance in validation metrics, while *β* = 0.05 converged smoothly (Figure 1B).

**Figure 1A.**
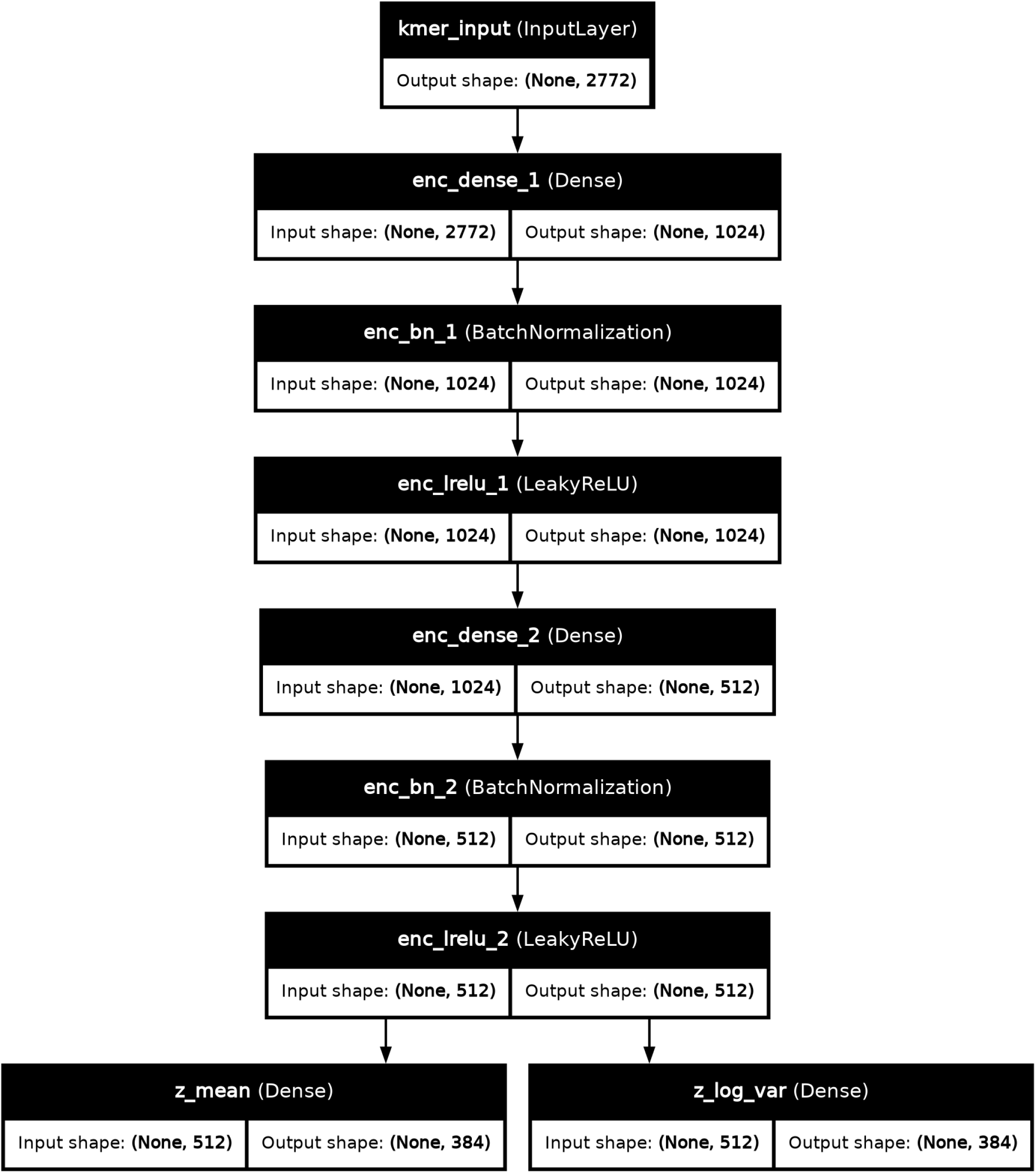
(left): Encoder architecture. Input: 2,772 CLR-transformed k-mer frequencies. Output: z_mean and z_log_var (384 dimensions each).

**Figure 1A:**
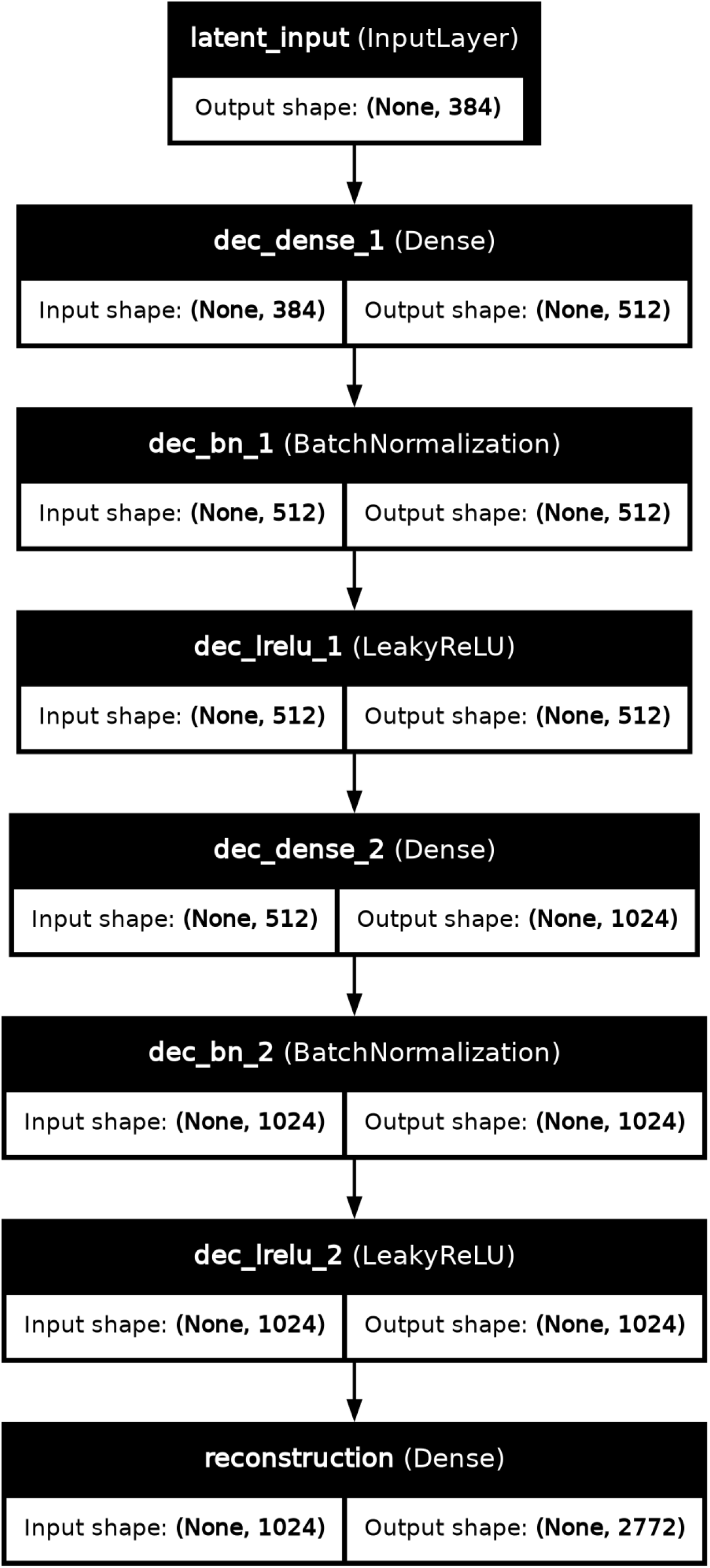
(right): Decoder architecture. Input: 384-dimensional latent vector. Output: 2,772 reconstructed k-mer frequencies.

**Figure 1B:**
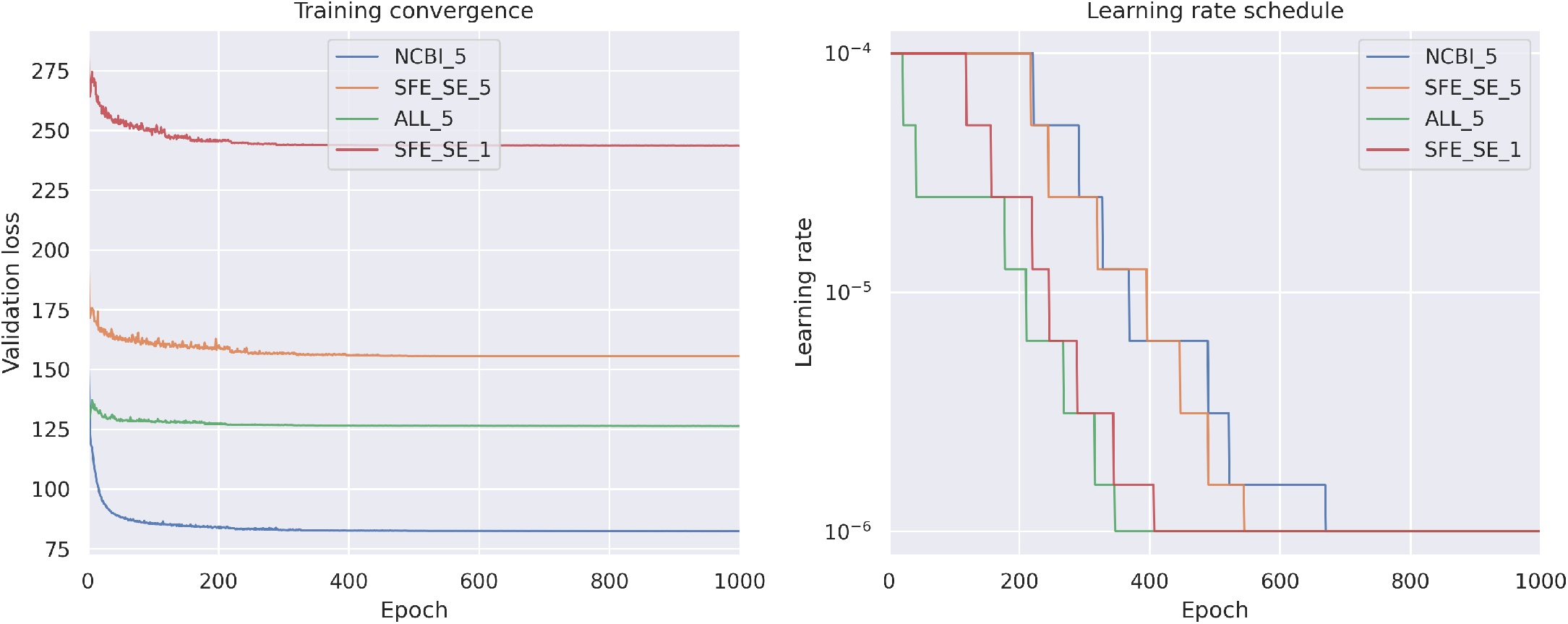
Training convergence (left) and learning rate schedule (right) for four key models.

The reconstruction error was dominated by 6-mers, which accounted for approximately 95% of the total MSE across all models, while 5-mers contributed ~4% and shorter k-mers (1–4) were reconstructed with high fidelity (Figure 1D). This distribution reflects the information bottleneck: compressing 2,080 hexamer frequencies into a 384-dimensional latent space is inherently lossy, whereas the smaller number of shorter k-mer features (10 dimers, 32 trimers, 136 tetramers) can be represented more accurately. The 6-mer dominance validates our decision to focus on k-mers up to length 6 rather than including 7-mers, as longer k-mers would likely exacerbate this bottleneck without proportional gains in discriminative power for our species-level clustering objective.

**Figure 1D:**
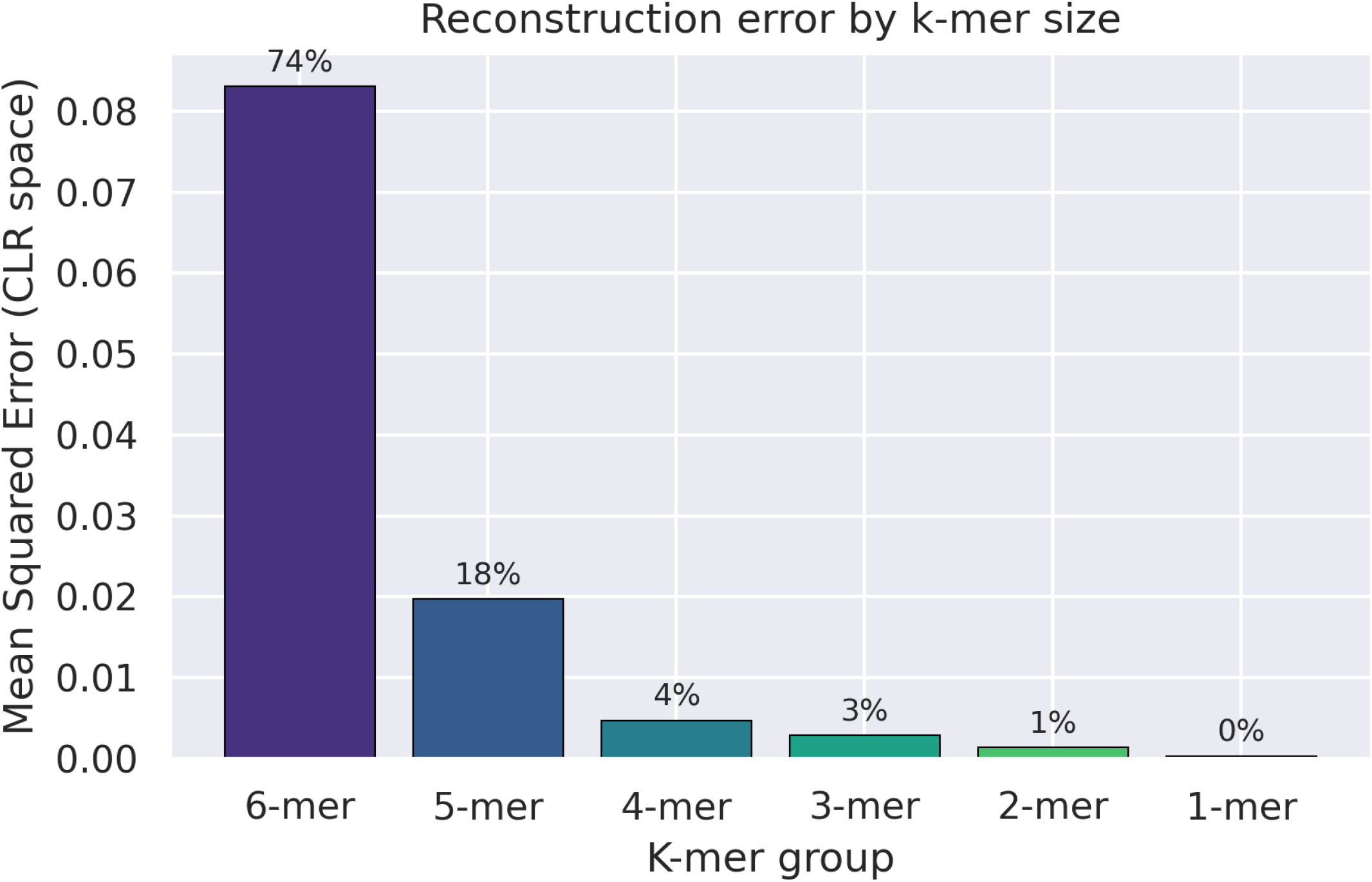
Per-k-mer reconstruction MSE breakdown showing 6-mers dominate the error

Based on this evaluation, we selected latent dimension = 384 and *β* = 0.05 as our final configuration (see Methods for details). This configuration was subsequently validated across all fifteen models (see below), confirming its suitability for downstream similarity search and clustering.

### Training Metrics Across Length Thresholds

We trained ten independent VAE models—five ALL (all four sources) and five brackish (SFE+SE only)— at minimum contig length thresholds of 1–5 kbp (Table 1 in Methods). All models used identical architecture and hyperparameters and were trained for 1,000 epochs on randomly shuffled data with a 90/10 train/validation split (Tables 2 and 3).

**Table 2:** Final training metrics for ALL models (epoch 1000).

| Min length | Contigs | Val loss | MSE | 6-mer MSE | KL | Train/Val gap |
| --- | --- | --- | --- | --- | --- | --- |
| 1 kbp | 17,415,045 | 189.18 | 0.059 | 0.077 | 476 | 0.12 |
| 2 kbp | 17,055,694 | 177.61 | 0.055 | 0.071 | 468 | 0.46 |
| 3 kbp | 16,495,774 | 165.93 | 0.052 | 0.068 | 462 | 0.33 |
| 4 kbp | 14,822,956 | 143.14 | 0.044 | 0.057 | 425 | 0.17 |
| 5 kbp | 13,439,298 | 126.43 | 0.038 | 0.049 | 395 | 0.26 |

**Table 3:** Final training metrics for brackish models (epoch 1000).

| Min length | Contigs | Val loss | MSE | 6-mer MSE | KL | Train/Val gap |
| --- | --- | --- | --- | --- | --- | --- |
| 1 kbp | 6,693,829 | 243.62 | 0.080 | 0.103 | 499 | 0.15 |
| 2 kbp | 6,455,140 | 224.24 | 0.073 | 0.095 | 492 | 0.07 |
| 3 kbp | 5,951,176 | 200.40 | 0.064 | 0.083 | 449 | -0.03 |
| 4 kbp | 5,309,235 | 174.75 | 0.056 | 0.072 | 414 | 0.13 |
| 5 kbp | 4,776,770 | 155.59 | 0.049 | 0.063 | 389 | -0.03 |

Validation loss decreased monotonically with longer minimum contig length in both model families (189$→126*forALL*, 244→$156 for brackish), reflecting both the reduced noise in k-mer frequency estimates for longer sequences and the smaller dataset sizes at higher thresholds. Train/validation gaps were negligible (<0.5 points) across all ten models.

6-mer features dominated the total MSE in both families (~95–98%). Brackish models showed 1.3–2.3× higher per-k-mer MSE than ALL models, with the gap widening for shorter k-mers (3-mers through 1-mers). Notably, the ALL 5 kbp model showed anomalously high 2-mer and 1-mer MSE compared to lower thresholds, attributable to underrepresentation of extreme high-GC organisms (>75% GC) that assemble into shorter contigs and are progressively lost at higher thresholds. The 3 kbp dataset retained four GC distribution peaks compared to three at 5 kbp, explaining the better reconstruction of extreme-GC sequences at the lower threshold.

Learning rate scheduling showed a clear difference between model families: ALL models triggered their first ReduceLROnPlateau reduction at epochs 21–22, while brackish models sustained learning at the initial rate much longer (epochs 118–217), suggesting a smoother loss surface for the less diverse brackish data. All models spent the majority of training (456–684 epochs) at the minimum learning rate of 10^−6^.

### Cross-Model Comparison Reveals That Training Data Properties Determine Embedding Quality

Because each model was trained on a different data distribution, comparing metrics on each model’s own data does not provide a fair assessment. We therefore evaluated all ten models—five ALL and five brackish— across multiple test datasets, producing cross-comparison matrices of Spearman rank correlations. We first examined the ALL models within their family (Table 4), then compared them against brackish models (Tables 5–6), and evaluated the brackish models on their own domain (Table 7).

**Table 4:**
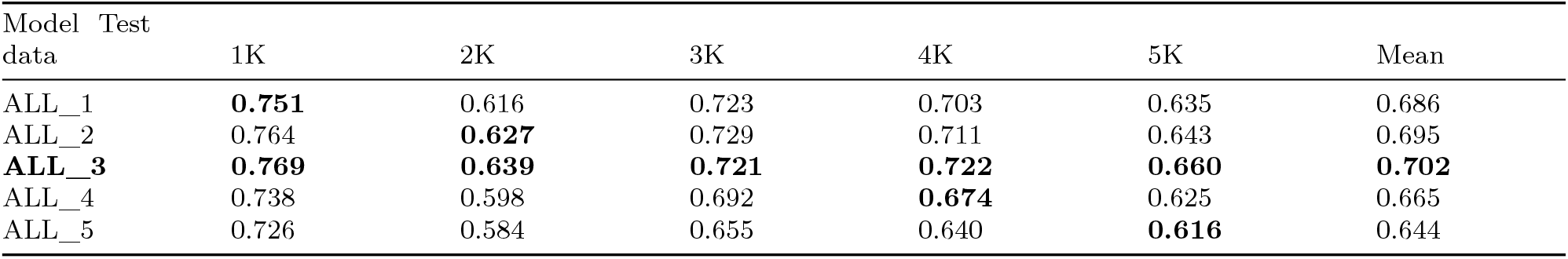
Spearman *ρ* for ALL models evaluated on all ALL evaluation datasets (held-out validation sets; see Methods). Bold diagonal entries are own-data results. Bold row indicates the best overall model (ALL_3), which achieved the highest Spearman *ρ* in four of five test datasets.

| Model Test data | 1K | 2K | 3K | 4K | 5K | Mean |
| --- | --- | --- | --- | --- | --- | --- |
| ALL_1 | **0.751** | 0.616 | 0.723 | 0.703 | 0.635 | 0.686 |
| ALL_2 | 0.764 | **0.627** | 0.729 | 0.711 | 0.643 | 0.695 |
| **ALL_3** | **0.769** | **0.639** | **0.721** | **0.722** | **0.660** | **0.702** |
| ALL_4 | 0.738 | 0.598 | 0.692 | **0.674** | 0.625 | 0.665 |
| ALL_5 | 0.726 | 0.584 | 0.655 | 0.640 | **0.616** | 0.644 |

**Table 5:** Brackish models evaluated on ALL test data (Spearman *ρ*).

| Model Test data | 1K | 2K | 3K | 4K | 5K | Mean |
| --- | --- | --- | --- | --- | --- | --- |
| **SFE_SE_1** | **0.907** | **0.820** | **0.881** | **0.843** | **0.829** | **0.856** |
| SFE_SE_2 | 0.889 | 0.808 | 0.873 | 0.827 | 0.807 | 0.841 |
| SFE_SE_3 | 0.886 | 0.802 | 0.868 | 0.813 | 0.811 | 0.836 |
| SFE_SE_4 | 0.878 | 0.804 | 0.850 | 0.807 | 0.805 | 0.829 |
| SFE_SE_5 | 0.866 | 0.790 | 0.836 | 0.787 | 0.782 | 0.812 |

**Table 6:** Grand comparison of all ten models on ALL test data.

| Model | Mean Spearman | Model type |
| --- | --- | --- |
| SFE_SE_1 | **0.856** | Brackish |
| SFE_SE_2 | 0.841 | Brackish |
| SFE_SE_3 | 0.836 | Brackish |
| SFE_SE_4 | 0.829 | Brackish |
| SFE_SE_5 | 0.812 | Brackish |
| ALL_3 | 0.702 | ALL |
| ALL_2 | 0.695 | ALL |
| ALL_1 | 0.686 | ALL |
| ALL_4 | 0.665 | ALL |
| ALL_5 | 0.644 | ALL |

**Table 7:** Brackish models evaluated on brackish test data (Spearman *ρ*).

| Model Test data | SFE_SE_1 | SFE_SE_2 | SFE_SE_3 | SFE_SE_4 | SFE_SE_5 | Mean |
| --- | --- | --- | --- | --- | --- | --- |
| **SFE_SE_1** | **0.773** | **0.739** | **0.723** | 0.816 | **0.779** | **0.766** |
| SFE_SE_2 | 0.743 | **0.714** | 0.682 | 0.784 | 0.758 | 0.736 |
| SFE_SE_3 | 0.749 | 0.704 | **0.676** | 0.776 | 0.736 | 0.728 |
| SFE_SE_4 | 0.751 | 0.684 | 0.687 | **0.781** | 0.726 | 0.726 |
| SFE_SE_5 | 0.715 | 0.674 | 0.643 | 0.736 | 0.691 | 0.692 |

ALL_3 was the best ALL model, achieving the highest mean Spearman *ρ* (0.702) and winning four of five columns. We observed a clear tier structure: ALL_3 (0.702) > ALL_1–2 (0.686–0.695) > ALL_4–5 (0.644– 0.665). Notably, no model was best on its own data—ALL_3 outperformed ALL_4 on 4K data and ALL_5 on 5K data—indicating that the 3 kbp threshold captures sufficient biological diversity (including extreme-GC organisms) to produce the most generalizable embeddings, even when evaluated on longer sequences.

The 2K test data proved uniquely challenging across all models (0.584–0.639), suggesting compositional heterogeneity specific to that length range. Shorter sequences also produced consistently higher nearest-neighbor MSE (0.16–0.18 for 1K/2K vs 0.10–0.14 for 3K–5K), reflecting the noisier k-mer profiles of short contigs.

Comparing across model families revealed a more striking pattern: brackish models (SFE+SE only) consistently outperformed ALL models on embedding quality—even when evaluated on ALL test data that included non-brackish and reference genome sequences not present in the brackish training data (Tables 5 and 6).

Every brackish model outperformed every ALL model (Figure 2D). The gap between the worst brackish model (SFE_SE_5, 0.812) and the best ALL model (ALL_3, 0.702) was +0.110 Spearman *ρ*. This demonstrates that training data properties—not quantity alone—determine embedding quality: the brackish models, with 4.8–6.7 million sequences from two sources, produced markedly better embeddings than the ALL models with 13.4–17.4 million sequences from four sources. Including the diverse non-brackish (FD) and reference genome (NCBI) data forced the model to spread its 384-dimensional latent space across a wider range of biology, diluting local distance structure for any particular data source.

**Figure 2A:**
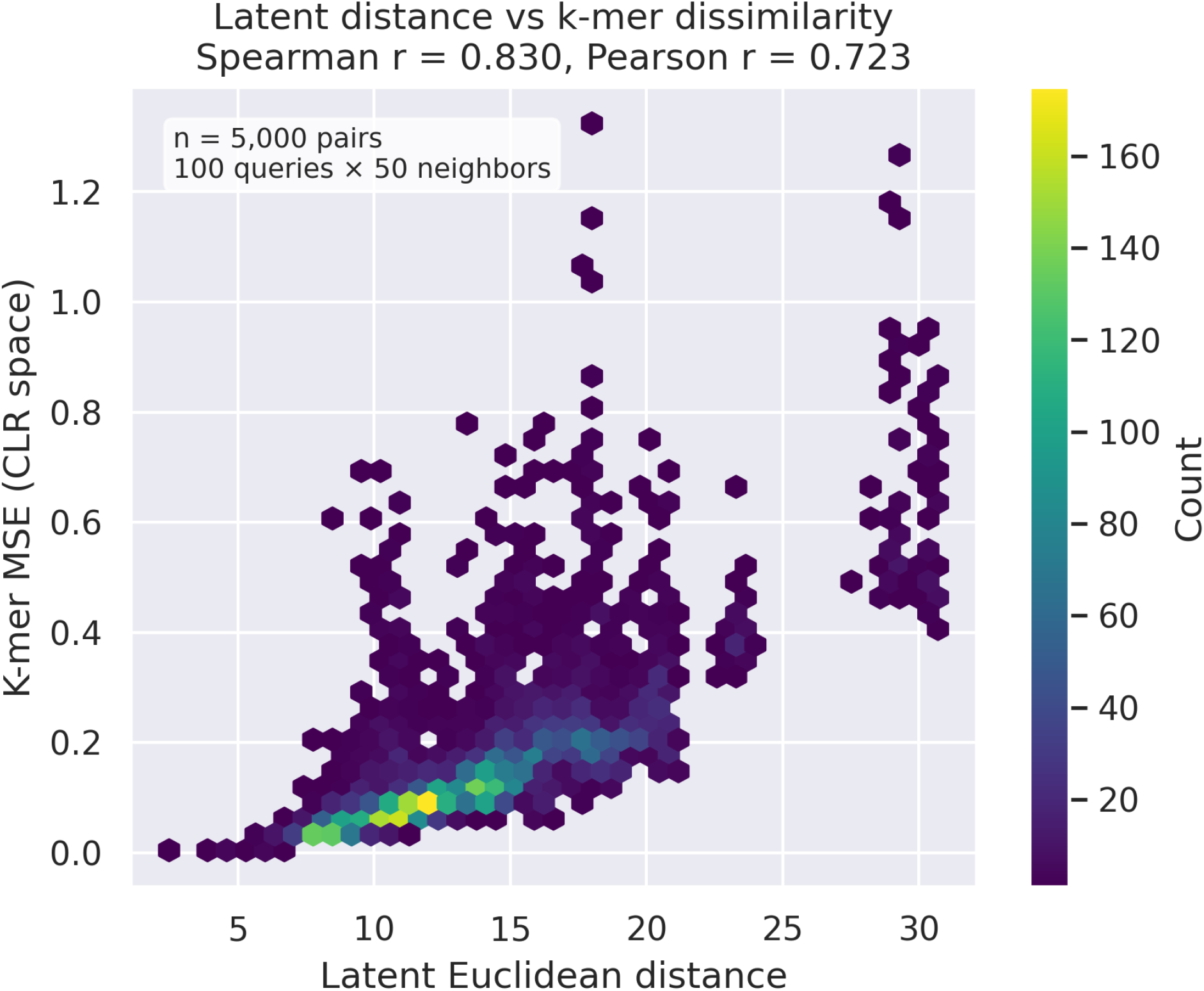
Scatter plot of latent Euclidean distance vs k-mer MSE for 50-nearest neighbors of 100 queries sampled from 50,000 sequences; hexbin or density plot to handle overplotting; show Spearman and Pearson correlations

**Figure 2B:**
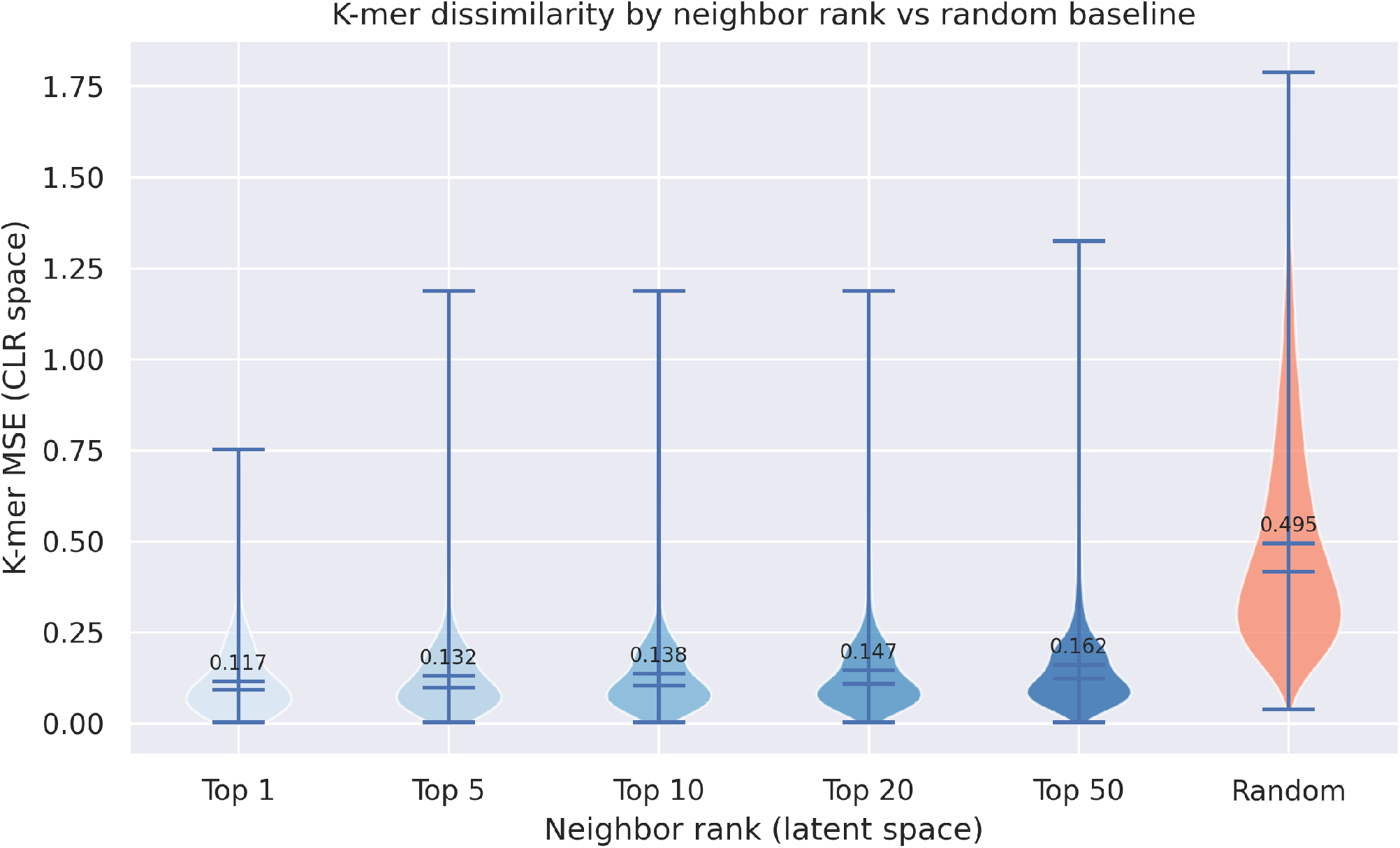
Box plots or violin plots showing distribution of k-mer MSE for different neighbor ranks compared to random baseline

When evaluated on brackish test data (Table 7; Figure 2C), the ranking was consistent: SFE_SE_1 achieved the highest mean Spearman of 0.766, with performance declining monotonically with increasing length threshold. This matches the augmented data ranking, confirming that lower thresholds—which include more diverse training sequences—consistently produce better-structured embeddings.

**Figure 2C:**
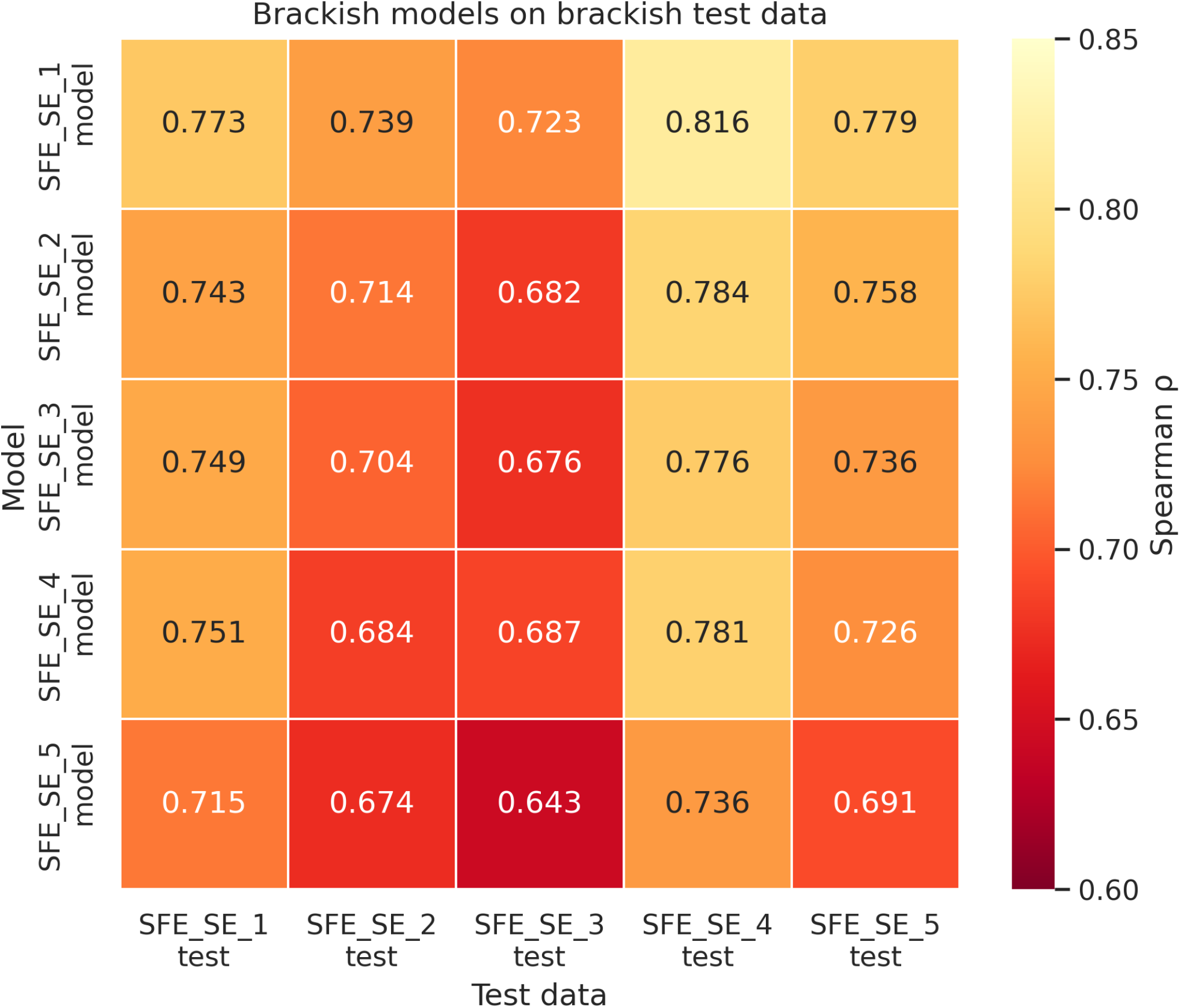
Spearman correlation heatmap for brackish models evaluated on brackish test data (5×5 cross-comparison).

**Figure 2D:**
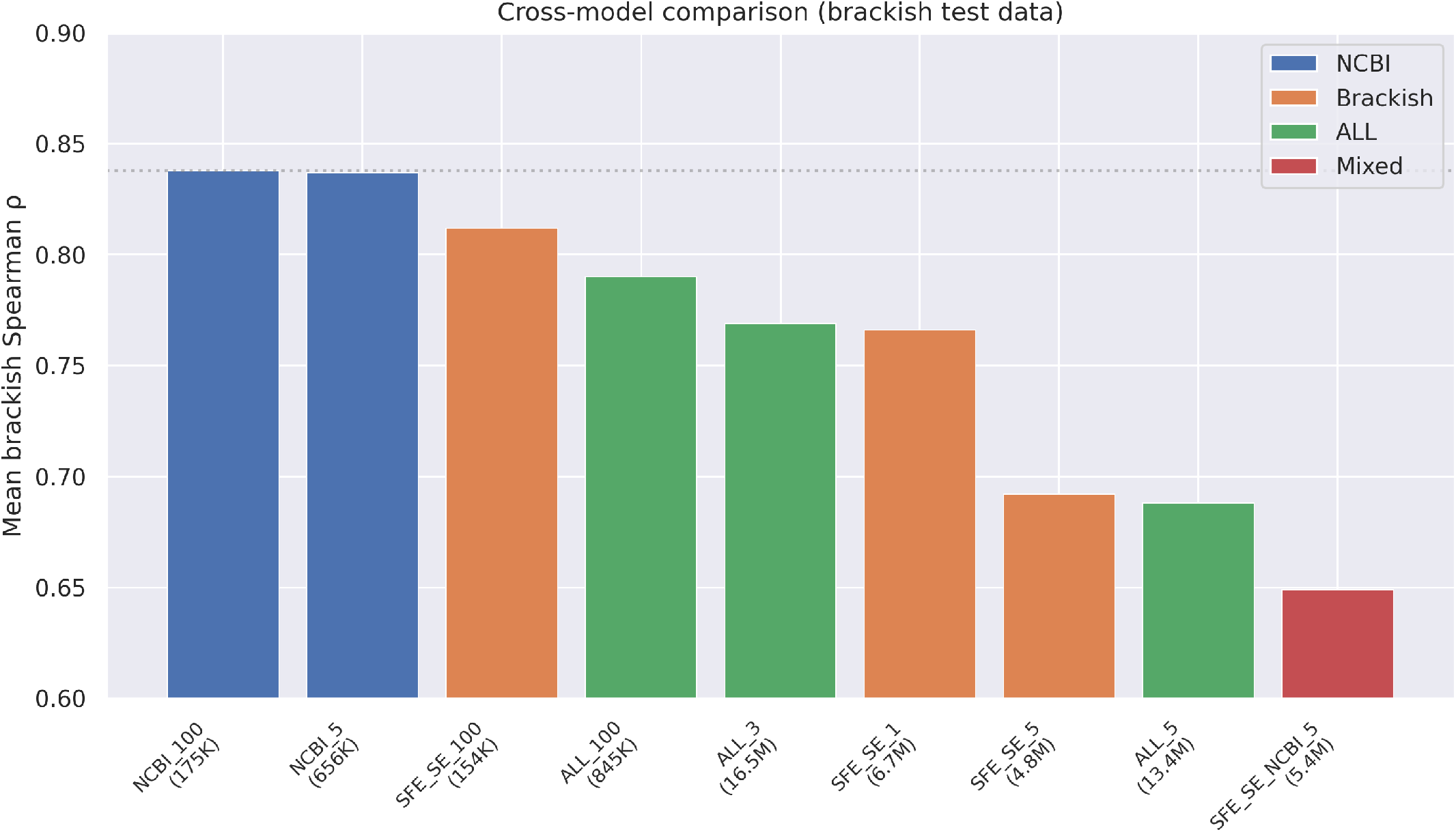
Mean brackish Spearman correlation across all nine models, colored by training data source.

However, five additional experimental models revealed that neither brackish training data nor Spearman correlation alone determine downstream clustering performance.

### Extended Comparison With Additional Models

The ten-model comparison established that lower length thresholds consistently produce better-structured embeddings. To probe the roles of training data source, length distribution, and dataset size, we trained five additional models (see Methods) and evaluated key models in an extended cross-comparison.

Several findings emerged from the extended comparison:

#### NCBI reference genome models dominate

Both NCBI_5 (0.837) and NCBI_100 (0.838) outperformed all brackish-trained models on brackish test data, despite never having seen environmental sequences during training. The best brackish model (SFE_SE_1, 0.766) fell 0.071 Spearman *ρ* below NCBI_5 despite being trained on 10× more sequences (6.7M vs 656K). This demonstrates that curated taxonomic diversity— ~20,000 species spanning Bacteria and Archaea—produces better-structured latent spaces than large volumes of domain-matched data.

#### NCBI reference genomes are readily organized

All models scored 0.89–0.95 on NCBI test data, including SFE_SE_5 which never saw reference genomes during training. This likely reflects the well-separated compositional signatures of curated reference genomes compared to the complex, overlapping compositions found in environmental metagenomes.

#### Training data properties matter more than source matching

On the evaluation domain that matters for clustering (SFE_SE_100, brackish contigs ≥ 100 kbp), NCBI_5 (Spearman 0.836) substantially outperformed SFE_SE_5 (0.766) despite never having seen brackish sequences. The NCBI and SFE_SE training data differ in several confounded ways: NCBI_5’s training data had a median contig length of ~37 kbp (vs. SFE_SE_5’s ~8 kbp), consists of curated complete genomes rather than fragmentary environmental contigs, and spans ~20,000 species across Bacteria and Archaea. The advantage cannot be attributed to a single factor—longer training contigs produce cleaner k-mer profiles, while genomic completeness ensures that each training example represents a coherent species-level composition. Filtering NCBI data to ≥ 100 kbp (NCBI_100) provided no additional benefit (0.832 vs 0.836), indicating that exact length matching is not required.

#### Mixing data sources can hurt

SFE_SE_NCBI_5 combined 4.8 million brackish sequences with 656,000 NCBI reference genomes—two seemingly complementary sources. Yet it achieved a mean Spearman correlation of only 0.649 on brackish data—the worst of any model. This demonstrates that naively combining training sources with different length distributions creates conflicting optimization targets that degrade embedding quality.

#### A small reference genome dataset outperforms domain-specific training

NCBI_5 (trained on just 656K contigs from ~20K reference genomes) outperformed SFE_SE_5 (trained on 4.8M domain-specific brackish contigs) on both Spearman correlation (0.837 vs 0.692 brackish mean) and downstream MCL clustering quality at 100 kbp, while achieving considerably better graph coverage (87% vs 80%; see below).

### Reconstruction Loss Does Not Predict Embedding Quality

A critical methodological finding emerged from comparing reconstruction MSE with embedding quality across models: the two metrics diverged across model families.

The ALL_5 model achieved the lowest reconstruction MSE (0.038, on its own ALL validation data) but the worst Spearman correlation (0.644) on common ALL test data (Figure 2E). Conversely, SFE_SE_5 had 1.3× higher reconstruction MSE (0.049, on its own brackish validation data) yet achieved 26% higher Spearman correlation (0.812 vs 0.644). The MSE values are not directly comparable because they are measured on different training distributions; the key comparison is the Spearman correlation, which is evaluated on the same ALL test data for all three models. This dissociation arises because reconstruction loss measures global fidelity of the input-output mapping, while embedding quality requires well-organized local geometry in the latent space. Spreading the latent space across four diverse sources (ALL) improves reconstruction by sharing features across data types, but dilutes the local distance structure needed for similarity retrieval.

**Figure 2E:**
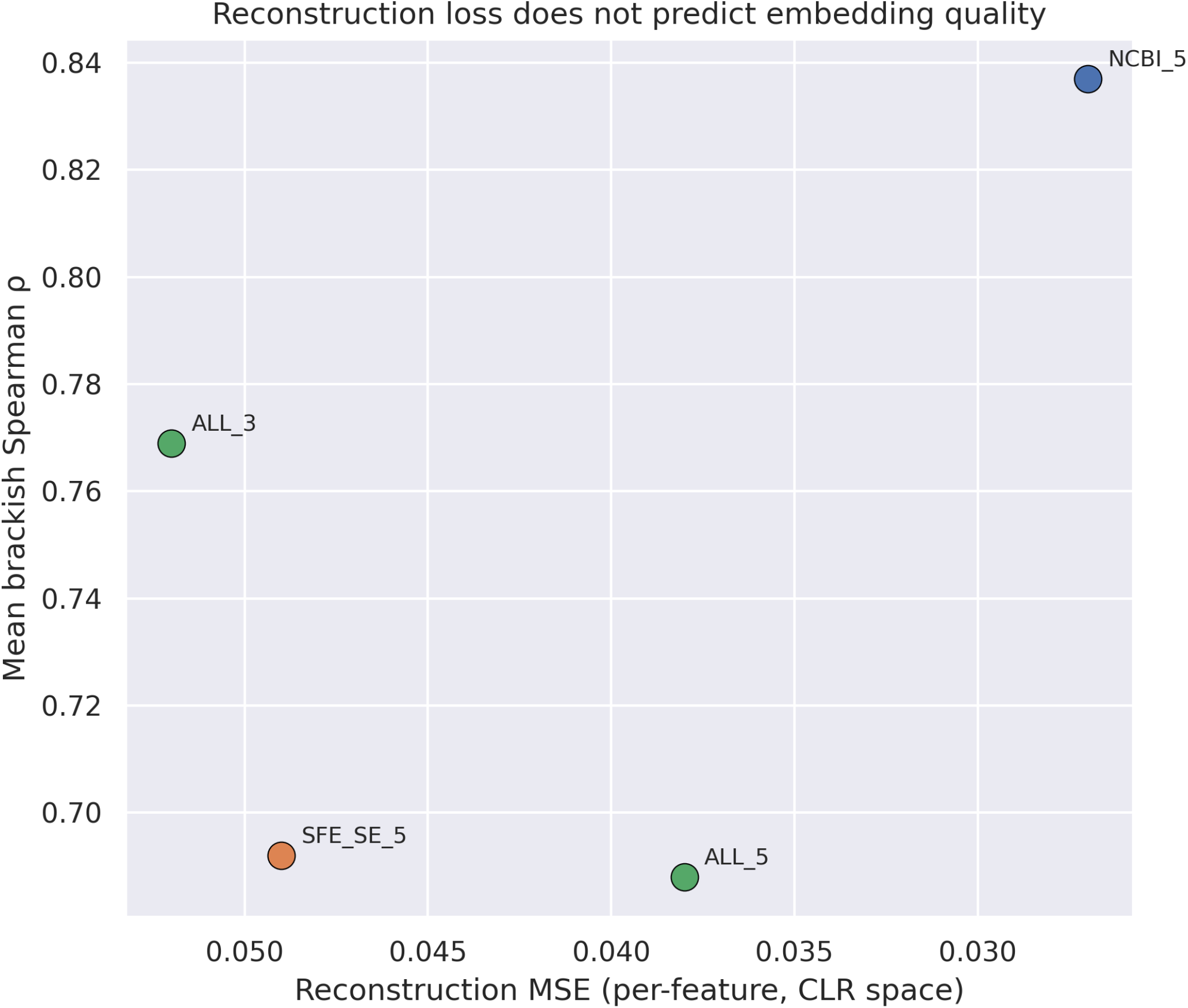
Reconstruction MSE vs Spearman correlation. Lower MSE does not predict better embedding quality.

Most published methods for metagenomic embedding report only reconstruction loss as a performance metric; these results demonstrate that an independent measure of local distance preservation is needed to evaluate whether an embedding is suitable for downstream clustering and retrieval. As shown below, even Spearman correlation itself does not reliably predict downstream task performance.

### The VAE Embedding Outperforms Linear Baselines on Taxonomic Coherence

To determine whether the VAE’s nonlinear compression provides value beyond simple dimensionality reduction, we compared VAE-384 against PCA-384 and raw CLR-2772 baselines through the identical clustering pipeline (see Methods).

The Spearman evaluation metric—which measures rank concordance between embedding distances and CLR-space MSE—is inherently circular for these baselines: raw CLR trivially achieves *ρ* = 1.000, PCA-384 achieves 0.948, and the VAE achieves 0.837. This confirms the finding from Table 9 that distance fidelity metrics do not predict clustering quality, since PCA preserves CLR distances better by construction—it is a linear projection of the very space the metric measures against.

GC content spans were tighter for PCA-384 (median 0.98 pp) than for the VAE (median 1.16 pp) at matched connectivity. However, PCA produced more clusters (11,717 vs 11,413 non-singleton), suggesting that tighter GC spans may reflect over-splitting rather than superior compositional coherence.

To resolve this ambiguity, we evaluated taxonomic coherence using GTDB-Tk direct classifications. The VAE outperformed both baselines on every combined metric at genus and species level (Table 10).

The advantage was most pronounced at genus level, where the VAE achieved 12% higher ARI and 17% higher pairwise F1 than PCA. At species level, all methods converged to similar V-measure (~0.94) and ARI (~0.67), though the VAE maintained a consistent edge. The advantage was driven by completeness rather than purity: PCA matched the VAE on cluster purity (both >99% at phylum through family) but fragmented taxa across more clusters.

Fragmentation analysis confirmed this pattern (Figure S2). Among species with ≥ 2 classified members, 77.9% were contained entirely within a single cluster for the VAE, compared to 66.4% for PCA—a gap of 11.5 percentage points (Table 11). Mean species-level completeness was 91.9% (VAE) vs 88.1% (PCA).

PCA’s tighter GC spans are thus an artifact of over-splitting: by distributing members of the same species across more clusters, each cluster is trivially more homogeneous in GC content but less complete taxonomically. The VAE’s learned nonlinear representation trades distance fidelity for a latent geometry in which related sequences cluster more completely—a property that does not emerge from linear dimensionality reduction. The relevant metric for evaluating embedding quality is therefore not distance preservation but downstream biological coherence.

### Euclidean Distance Outperforms Cosine in the Latent Space

We compared Euclidean and cosine distance metrics for nearest-neighbor retrieval in the latent space. Using the ALL_4 model evaluated on its own 4K held-out data, Euclidean distance achieved Spearman *ρ* = 0.697, compared to 0.621 for cosine distance—an improvement of +0.076. This advantage is expected because the decoder learns a smooth mapping from latent to output space, and the MSE loss penalizes squared L2 distance in the reconstruction space; consequently, points close in Euclidean latent distance tend to produce similar reconstructions, making Euclidean distance a natural proxy for compositional similarity.

Cosine distance, which measures the angle between vectors rather than their absolute separation, showed poor discriminative range: in 384 dimensions, the standard deviation of pairwise cosine distances was only 0.061— insufficient for meaningful nearest-neighbor ranking. While both metrics can in principle suffer from the concentration of distances phenomenon^1^ in high-dimensional spaces, Euclidean distance retained sufficient discriminative range in this latent space. All subsequent analyses used Euclidean distance.

### Local Distance Preservation Enables Reliable Similarity Search

A fundamental requirement for our embedding is that sequences with similar k-mer composition should map to nearby points in latent space, as clustering and nearest-neighbor retrieval depend on the reliability of distance comparisons. We rigorously validated this property by testing whether latent-space Euclidean distances accurately predict k-mer dissimilarity (MSE between CLR-transformed k-mer frequency vectors) in the original high-dimensional space (see Methods for protocol details).

Across all models and evaluation conditions, we observed strong monotonic relationships between latent distance and k-mer dissimilarity. On the ALL cross-comparison, the best model (ALL_3) achieved Spearman *ρ* = 0.639–0.769 across test datasets, with a mean of 0.702. The NCBI reference genome models achieved the highest correlations on brackish data (NCBI_5 mean *ρ* = 0.837, range 0.817–0.856 across five brackish test datasets). Spearman correlations were consistently higher than Pearson correlations, indicating a monotonic but nonlinear relationship between latent distance and k-mer MSE—a property well-suited for ranking-based retrieval where only the ordering of neighbors matters.

To quantify the practical benefit of the embedding, we compared the k-mer dissimilarity of nearest neighbors in latent space to random baseline pairs. Across all ALL models on the common test set, the closest neighbor (rank 1) had a mean k-mer MSE of 0.100–0.134 depending on the model, compared to the random baseline of ~0.5—a 3.7–5.0× improvement. K-mer MSE increased gradually from top-1 to top-50 neighbors, demonstrating that distance ranking in latent space accurately reflects compositional similarity. This property is essential for k-nearest-neighbor queries and for graph-based clustering approaches that rely on local neighborhood structure.

### Latent Space Structure: An Archipelago of Species

#### GC content as the primary axis of variation

Visualization of the latent space using t-SNE^2^ (openTSNE 1.0.4, default perplexity = 30, early exaggeration = 12) on the 4.8 million SFE_SE_5 embeddings revealed that GC content is the dominant axis of variation, with two major lobes corresponding to low-GC (20–40%) and high-GC (50–70%) organisms. This structure reflects the fundamental role of base composition in shaping k-mer frequencies^3^: GC content determines 1-mer frequencies and strongly influences higher-order k-mer patterns through codon usage bias and dinucleotide preferences. Sequences from different datasets and environments (SFE, SE) intermixed throughout the space rather than forming source-specific clusters, indicating that the VAE learned generalizable sequence patterns rather than memorizing dataset-specific features.

#### Archipelago structure

The distribution of sequences in the latent space is discrete, not continuous. Each species or lineage occupies a tiny, dense island surrounded by empty space—an archipelago of compositionally distinct groups rather than a continuum of gradual transitions. This structure is quantified by the nearest-neighbor distance analysis (Table 12): at 10 kbp minimum length, only 13.8% of sequences have a nearest neighbor within the clustering threshold of Euclidean distance 5 (median nn1 = 9.20); this pattern is consistent across models. The transition from isolated to connected is nearly a step function: for any given sequence, either all nearest neighbors cluster tightly within a small radius (dense island) or the nearest neighbor is distant (isolated singleton), with few intermediate cases.

This step-function behavior (Figure 3B) reflects the compositional discreteness of natural microbial communities. Each species or lineage occupies a narrow region of k-mer space, and the latent space preserves these compositional gaps rather than interpolating between them. Lineages sampled deeply enough to produce multiple assembled contigs form dense islands, while those contributing only a handful of contigs (or even just one) appear as isolated singletons with no nearby neighbors. Assembly fragmentation also contributes: larger genomes fragment into more contigs above the length threshold, producing denser clusters than their species count alone would predict.

**Figure 3A:**
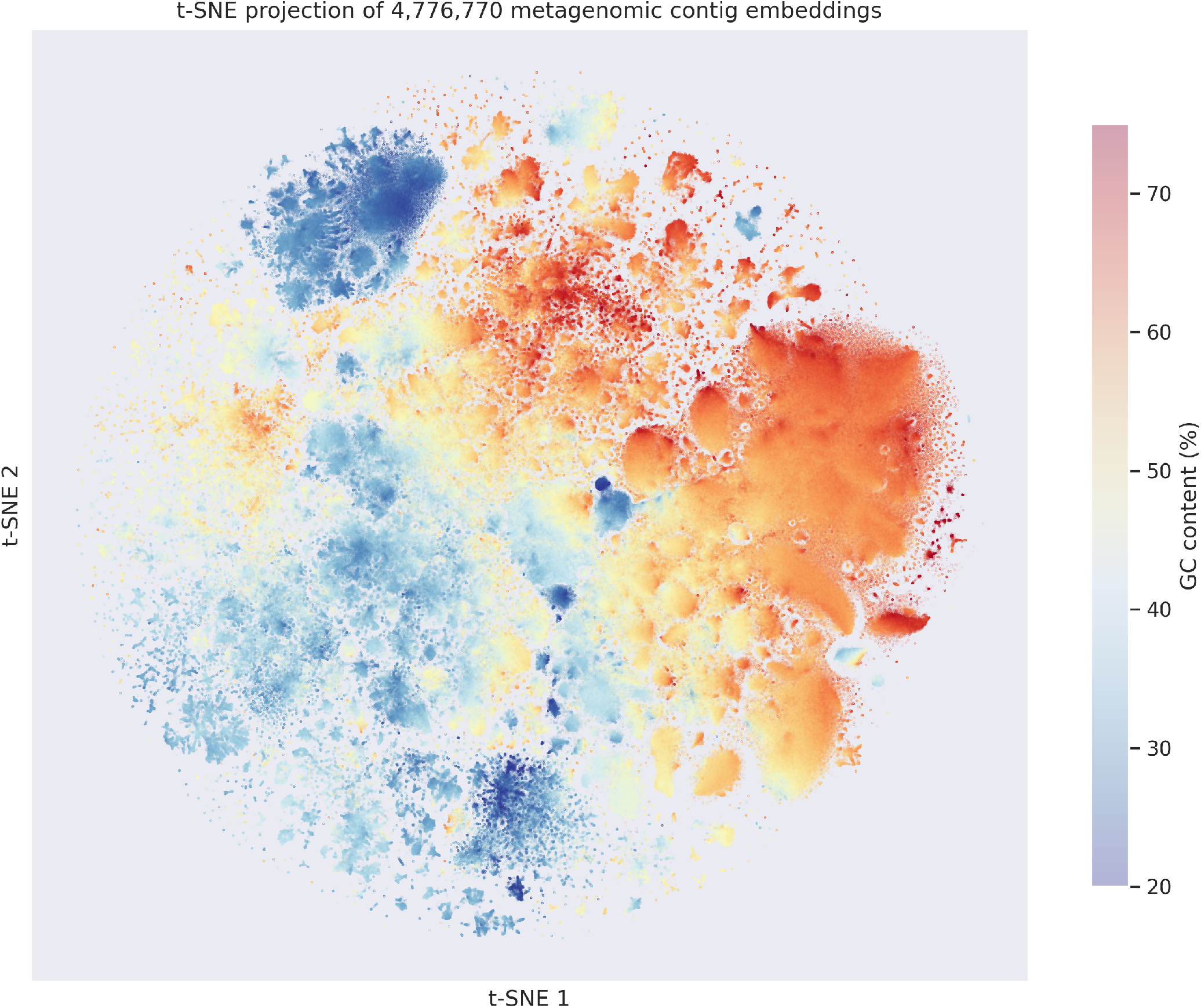
t-SNE projection of 4.8M embeddings colored by GC content (continuous scale); two lobes visible

**Figure 3B:**
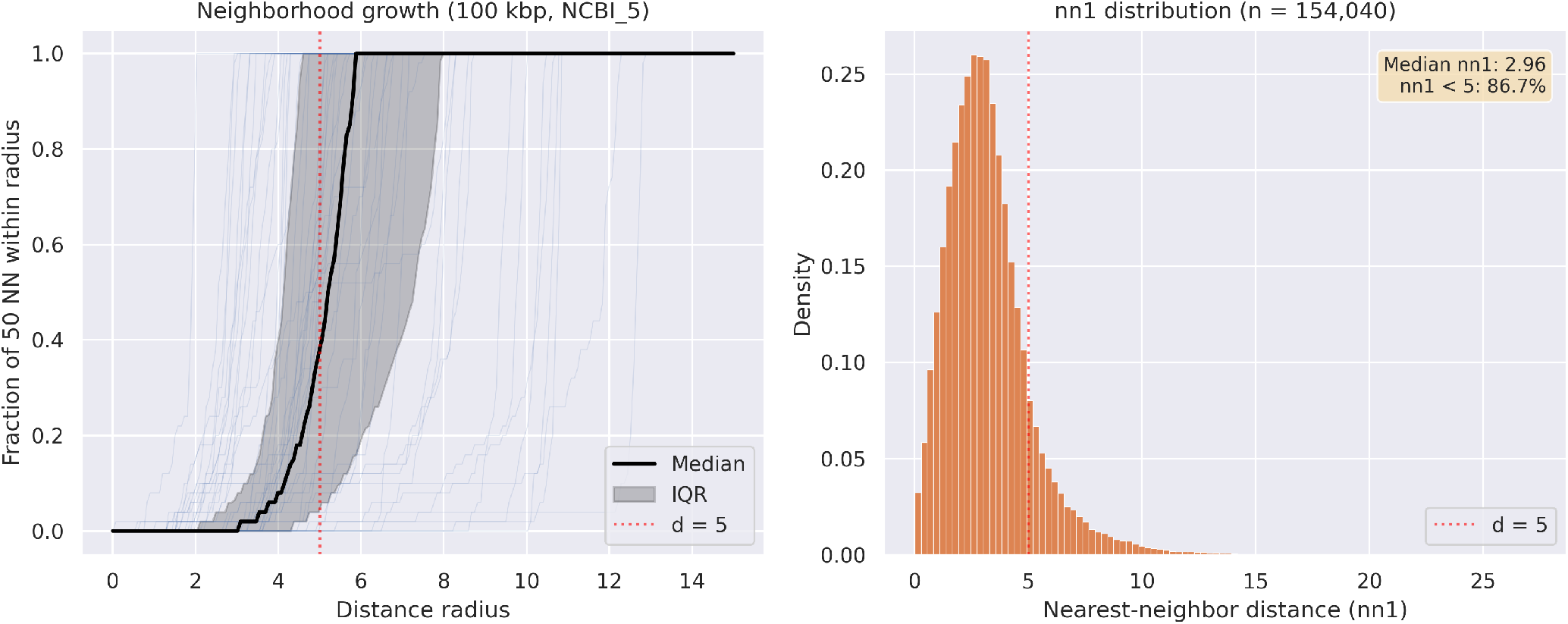
Neighborhood growth (left) showing step-function shape characteristic of archipelago structure. nn1 distribution (right) at 100 kbp.

#### Intrinsic dimensionality

Despite the 384-dimensional ambient space, the data lies on a low-dimensional manifold. The TWO-NN estimator^4^ yielded intrinsic dimensionality estimates that decreased with increasing minimum contig length.

The decreasing dimensionality (Figure 3C) reflects the loss of noise dimensions as shorter, noisier contigs are filtered out. Because TWO-NN estimates dimensionality from local neighbor-distance ratios, these values characterize the within-island manifold structure rather than the global arrangement of the archipelago. At 100 kbp, the effective dimensionality of 3.74 suggests that just four dimensions capture the essential variation among long contigs—consistent with GC content plus a small number of additional compositional axes (e.g., dinucleotide biases, codon usage). The neighbor count within radius r grew as a low-order power of r (R^2^ = 0.954 for a power-law fit), supporting the interpretation that the data lie on a low-dimensional manifold rather than filling the 384-dimensional ambient space.

**Figure 3C:**
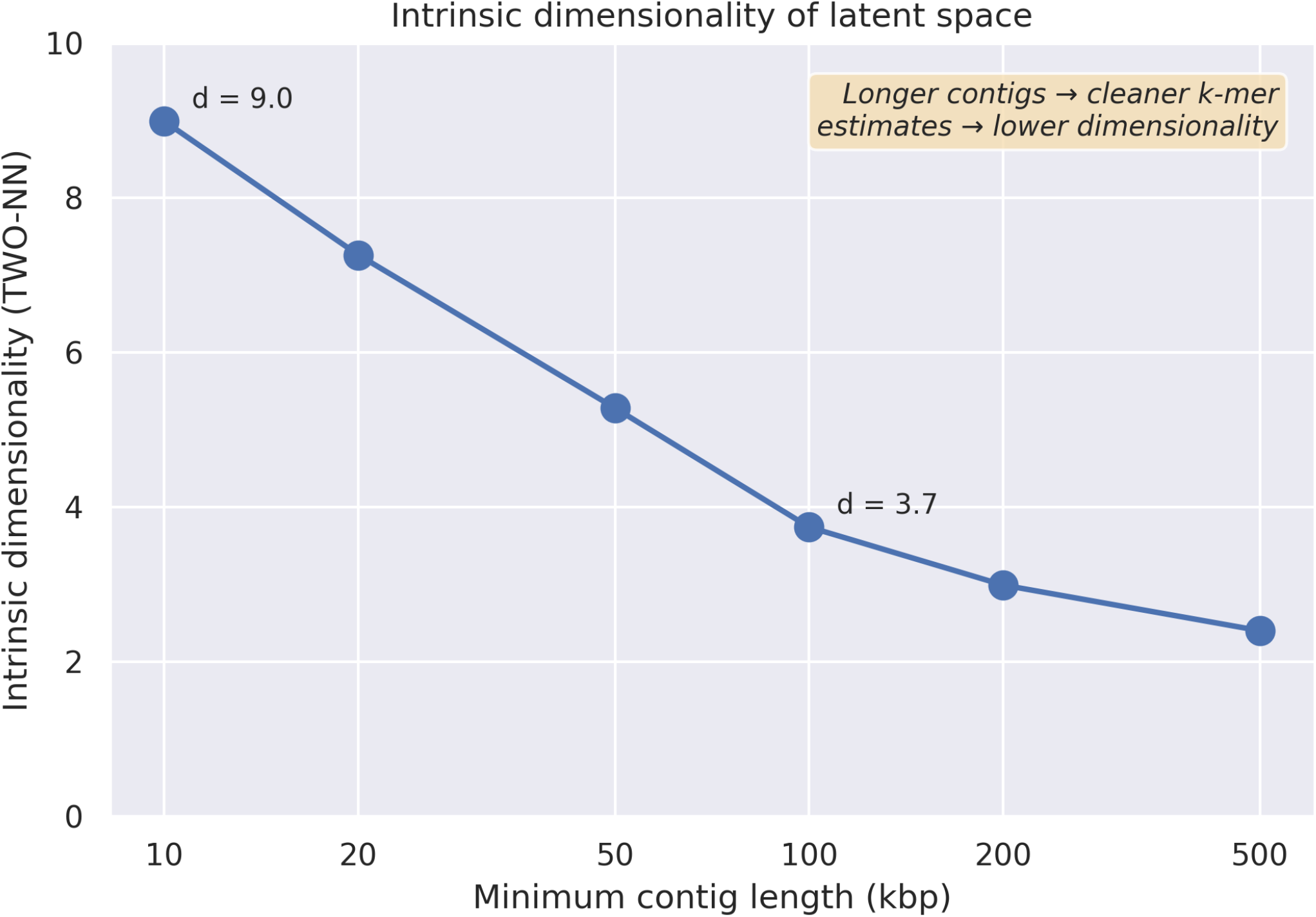
Intrinsic dimensionality (TWO-NN) decreases from ~9 at 10 kbp to ~4 at 100 kbp.

### Hub Nodes Cause Giant Component Formation

Analysis of the directed k-NN graph (each sequence points to its k = 50 nearest neighbors) revealed a severe hubness problem^5,6^. On the 10 kbp brackish dataset (3.0 million contigs, SFE_SE_5 model, d = 10), in-degree (the number of times a sequence appears in other sequences’ neighbor lists) was extremely right-skewed: the top hub had an in-degree of 58,377—2% of the entire dataset pointing to a single sequence. The top 20 hubs were all large brackish contigs (400 kbp–1.2 Mbp) with ~28–29% GC content, occupying a region of the latent space where many sequences’ nearest-neighbor lists converge.

These hubs cause a critical problem for graph-based clustering (Figure 4A). In the symmetric k-NN graph, hubs act as transitivity chain anchors: if sequences A and B both include the same hub in their neighbor lists, the symmetric graph connects A–hub–B, even if A and B are compositionally dissimilar. At a distance threshold of d = 10 on 10 kbp data, this chaining effect produced a giant connected component containing 1,297,780 sequences (77% of all clustered sequences), with GC content ranging from 20% to 82%—a span too wide to represent any compositionally coherent group.

**Figure 4A:**
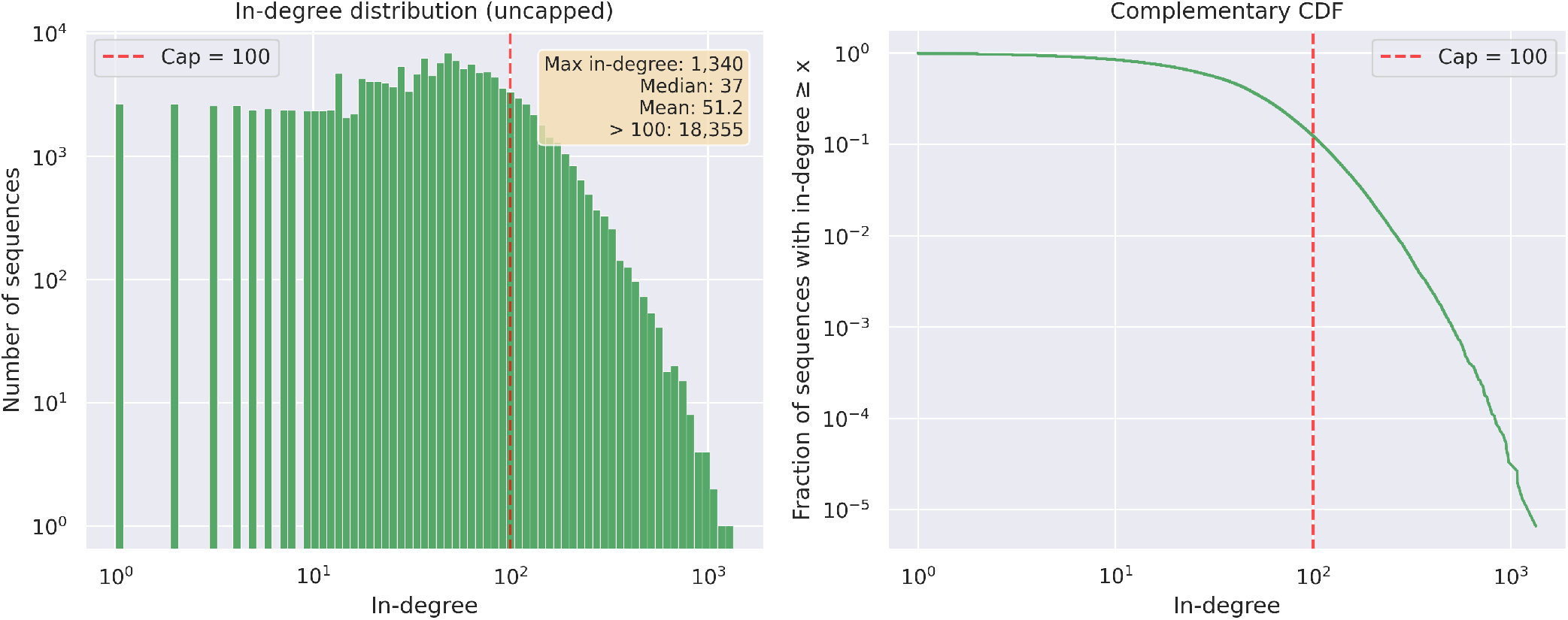
In-degree distribution (left) and complementary CDF (right) showing hub nodes in the uncapped k-NN graph. Red line: cap = 100.

**Figure 4B:**
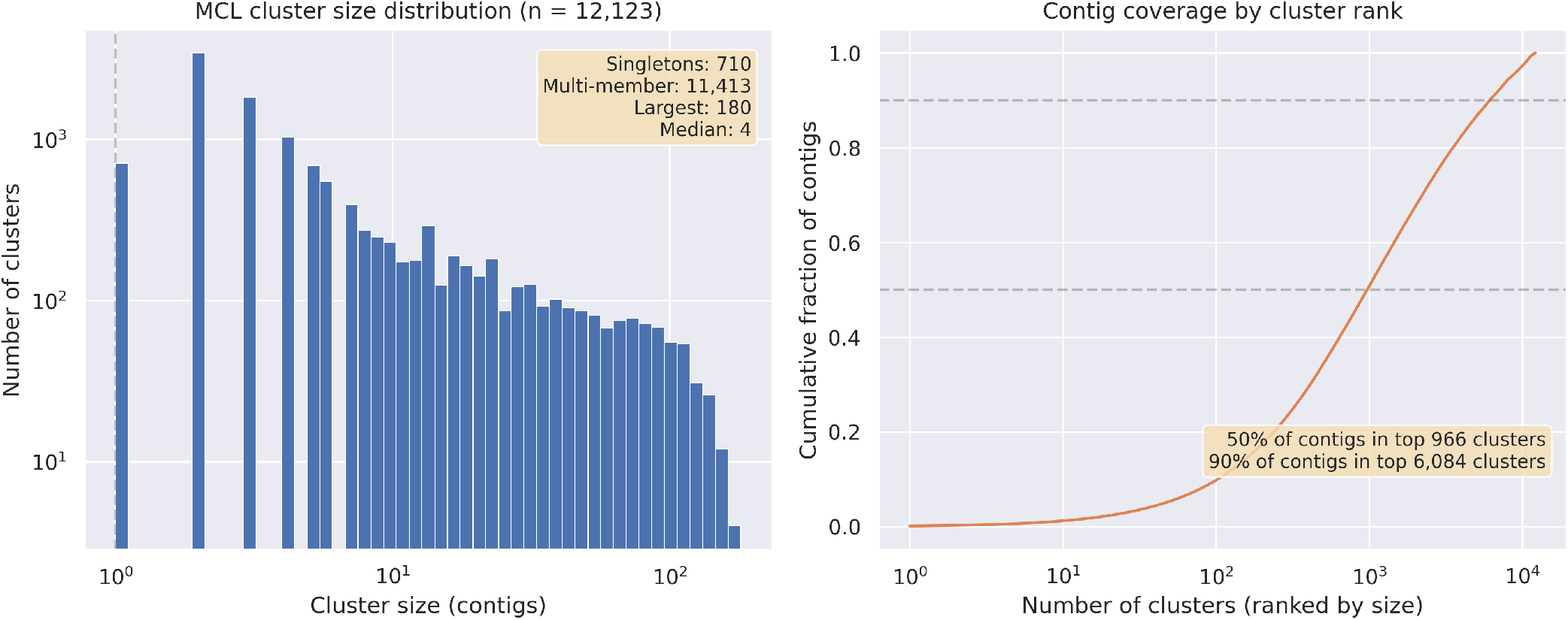
MCL cluster size distribution (left) and cumulative contig coverage by cluster rank (right). 12,123 clusters at 100 kbp, d = 5, I = 3.0.

We tested three graph construction strategies (see Methods). Mutual k-NN filtering eliminated hubs entirely but raised the singleton rate from 51% to 85%, severely reducing coverage. In-degree capping at 100 provided the best compromise: it reduced the largest community from 119,000 to 56,000 sequences while maintaining ~47% coverage at 10 kbp d = 10.

### MCL Outperforms Modularity-Based Clustering

MCL substantially outperformed Leiden on metagenome nearest-neighbor graphs where hub nodes arise. On the same in-degree-capped graph at 10 kbp with distance threshold d = 10 (SFE_SE_5 model), the GC content spans of the three largest communities were:

| Method | C1 (pp) | C2 (pp) | C3 (pp) |
| --- | --- | --- | --- |
| Leiden (resolution 1.0) | 33 | 49 | 42 |
| MCL (I = 3.0) | 9 | 7 | 7 |
| MCL (I = 5.0) | 9 | 7 | 6 |

MCL achieved GC spans 3.7–7× tighter than Leiden at the same configuration (Figure 4C). This advantage reflects the difference in how each algorithm handles hub-mediated connections: MCL’s flow simulation progressively weakens indirect paths through hubs (see Methods), while Leiden’s modularity optimization tends to merge compositionally dissimilar nodes connected through hubs when the hub-mediated edge density exceeds the null model expectation.

**Figure 4C:**
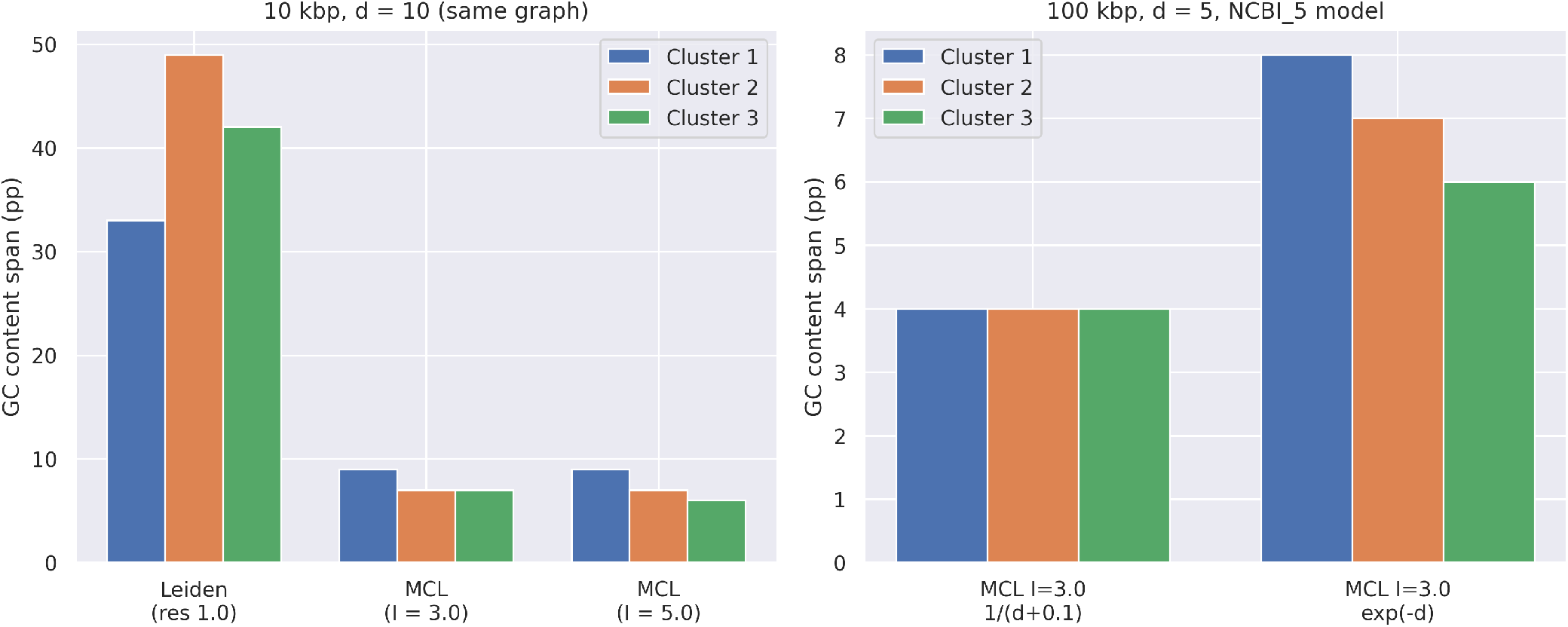
GC content span comparison. Left: MCL vs Leiden on same graph (10 kbp). Right: weight function comparison at 100 kbp.

#### Weight function comparison

The choice of edge weight function substantially affected MCL performance. At 100 kbp with distance threshold d = 7 (SFE_SE_5 model):

| Weight function | I = 1.4 GC spans | I = 3.0 GC spans |
| --- | --- | --- |
| $1/(d + 0.1)$ | 13, 12, 18 pp | 4, 5, 5 pp |
| $\exp(-d)$ | 21, 18, 14 pp | 8, 7, 6 pp |

The gentle 1/(d + 0.1) weighting outperformed exponential weighting at every inflation value. This counter-intuitive result occurs because MCL needs flow to traverse the local neighborhood before inflation cuts weak connections: exp(–d) over-suppresses medium-range edges (d = 3–5) that MCL uses for cluster boundary definition, starving the expansion step of the flow it needs to identify community structure.

### Sequence Length Drives Clustering Quality

The preceding analyses used the SFE_SE_5 model, which was the initial candidate for downstream clustering based on early cross-comparison results; the structural findings about hub nodes, MCL vs Leiden, and weight functions are properties of the graph topology and hold across models. Because NCBI_5 outperformed all brackish models on both full brackish data (Spearman 0.837 vs 0.766 for the best brackish model SFE_SE_1; Table 8) and on ≥ 100 kbp contigs (0.836 vs 0.766), we switched to the NCBI_5 model for the remaining analyses. We computed the nearest-neighbor distance (nn1) for 10,000 random queries at each length threshold, searching against all sequences at that threshold.

**Table 8:**
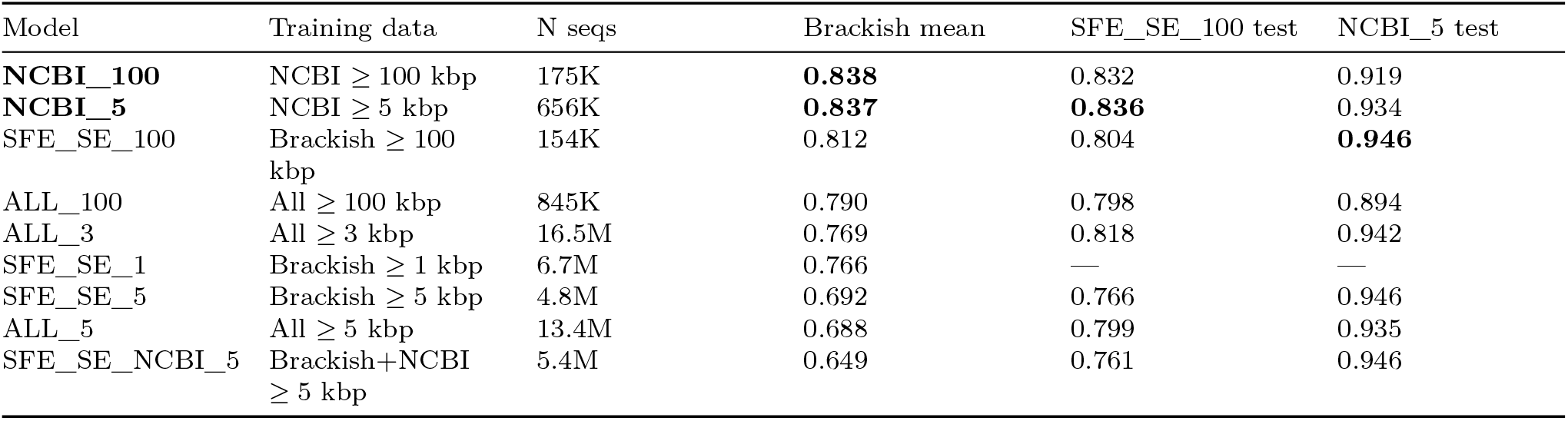
Extended cross-model comparison (Spearman *ρ*). “Brackish mean” is the mean Spearman across all five brackish test datasets at standardized 50K pool size. “SFE_SE_100 test” and “NCBI_5 test” use the respective held-out validation sets. Models ranked by brackish mean.

| Model | Training data | N seqs | Brackish mean | SFE_SE_100 test | NCBI_5 test |
| --- | --- | --- | --- | --- | --- |
| **NCBI_100** | NCBI $\geq 100$ kbp | 175K | **0.838** | 0.832 | 0.919 |
| **NCBI_5** | NCBI $\geq 5$ kbp | 656K | **0.837** | **0.836** | 0.934 |
| SFE_SE_100 | Brackish $\geq 100$ kbp | 154K | 0.812 | 0.804 | **0.946** |
| ALL_100 | All $\geq 100$ kbp | 845K | 0.790 | 0.798 | 0.894 |
| ALL_3 | All $\geq 3$ kbp | 16.5M | 0.769 | 0.818 | 0.942 |
| SFE_SE_1 | Brackish $\geq 1$ kbp | 6.7M | 0.766 | — | — |
| SFE_SE_5 | Brackish $\geq 5$ kbp | 4.8M | 0.692 | 0.766 | 0.946 |
| ALL_5 | All $\geq 5$ kbp | 13.4M | 0.688 | 0.799 | 0.935 |
| SFE_SE_NCBI_5 | Brackish+NCBI $\geq 5$ kbp | 5.4M | 0.649 | 0.761 | 0.946 |

**Table 9:** Reconstruction loss vs embedding quality. Spearman *ρ* is the mean across the five ALL test datasets (Table 6), providing a fair comparison on common evaluation data.

| Model | Training MSE | Spearman $\rho$ (ALL test data) | Relationship |
| --- | --- | --- | --- |
| ALL_5 | **0.038** (lowest) | 0.644 (worst) | Best reconstruction, worst embedding |
| ALL_3 | 0.052 | 0.702 |  |
| SFE_SE_5 | 0.049 | **0.812** (best) | 1.3× higher MSE, 26% better Spearman |

**Table 10:** Taxonomic coherence of VAE, PCA, and CLR clustering. All methods evaluated on the same 26,841 GTDB-Tk classified contigs at ≥ 100 kbp.

| Metric | VAE-384 | PCA-384 | CLR-2772 |
| --- | --- | --- | --- |
| V-measure (genus) | 0.882 | 0.874 | 0.879 |
| V-measure (species) | 0.941 | 0.940 | 0.938 |
| ARI (genus) | 0.228 | 0.202 | 0.218 |
| ARI (species) | 0.669 | 0.668 | 0.661 |
| Pairwise F1 (genus) | 0.228 | 0.195 | 0.219 |
| Pairwise F1 (species) | 0.667 | 0.666 | 0.644 |

**Table 11:** Species completeness and fragmentation. Completeness measures the fraction of each taxon’s classified contigs in the largest cluster. “Perfect” indicates all classified contigs of that taxon are in a single cluster.

| Metric | VAE-384 | PCA-384 | CLR-2772 |
| --- | --- | --- | --- |
| Mean completeness (species) | 91.9% | 88.1% | 90.1% |
| Perfect completeness (species) | 77.9% | 66.4% | 73.0% |
| Mean completeness (genus) | 80.7% | 76.7% | 80.0% |
| Perfect completeness (genus) | 55.1% | 46.3% | 52.5% |

**Table 12:** Nearest-neighbor distance by minimum contig length.

| Threshold | Sequences | Mean nn1 | Median nn1 | nn1 < 5 (%) |
| --- | --- | --- | --- | --- |
| 3 kbp | 5,951,176 | 12.76 | 12.28 | 9.5 |
| 5 kbp | 4,776,770 | 11.43 | 11.17 | 10.0 |
| 10 kbp | 3,039,927 | 9.27 | 9.20 | 13.8 |
| 20 kbp | 1,556,556 | 7.11 | 7.02 | 24.8 |
| **50 kbp** | **461,674** | **4.99** | **4.82** | **53.4** |
| **100 kbp** | **154,041** | **3.58** | **3.38** | **81.6** |
| 200 kbp | 53,138 | 2.64 | 2.36 | 92.8 |
| 500 kbp | 11,006 | 2.08 | 1.70 | 93.4 |

**Table 12b:**
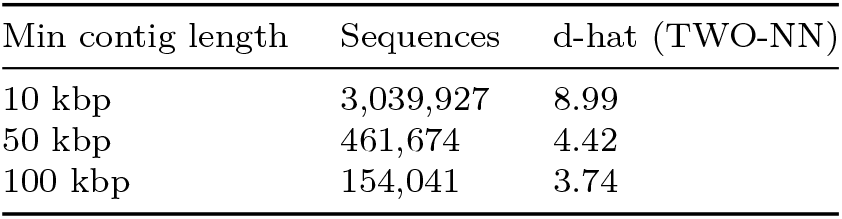
Intrinsic dimensionality by minimum contig length (TWO-NN estimator).

| Min contig length | Sequences | d-hat (TWO-NN) |
| --- | --- | --- |
| 10 kbp | 3,039,927 | 8.99 |
| 50 kbp | 461,674 | 4.42 |
| 100 kbp | 154,041 | 3.74 |

The nn1 distribution reveals a clear knee at 50–100 kbp (Figure 5A). Below 50 kbp, the majority of sequences are noise-dominated singletons (median nn1 > 5), while above 200 kbp, diminishing returns set in with a rapidly shrinking dataset. The 100 kbp threshold represents the optimal trade-off: 81.6% of sequences have a nearest neighbor within d = 5, and 154,041 contigs provide ample material for community detection.

**Figure 5A:**
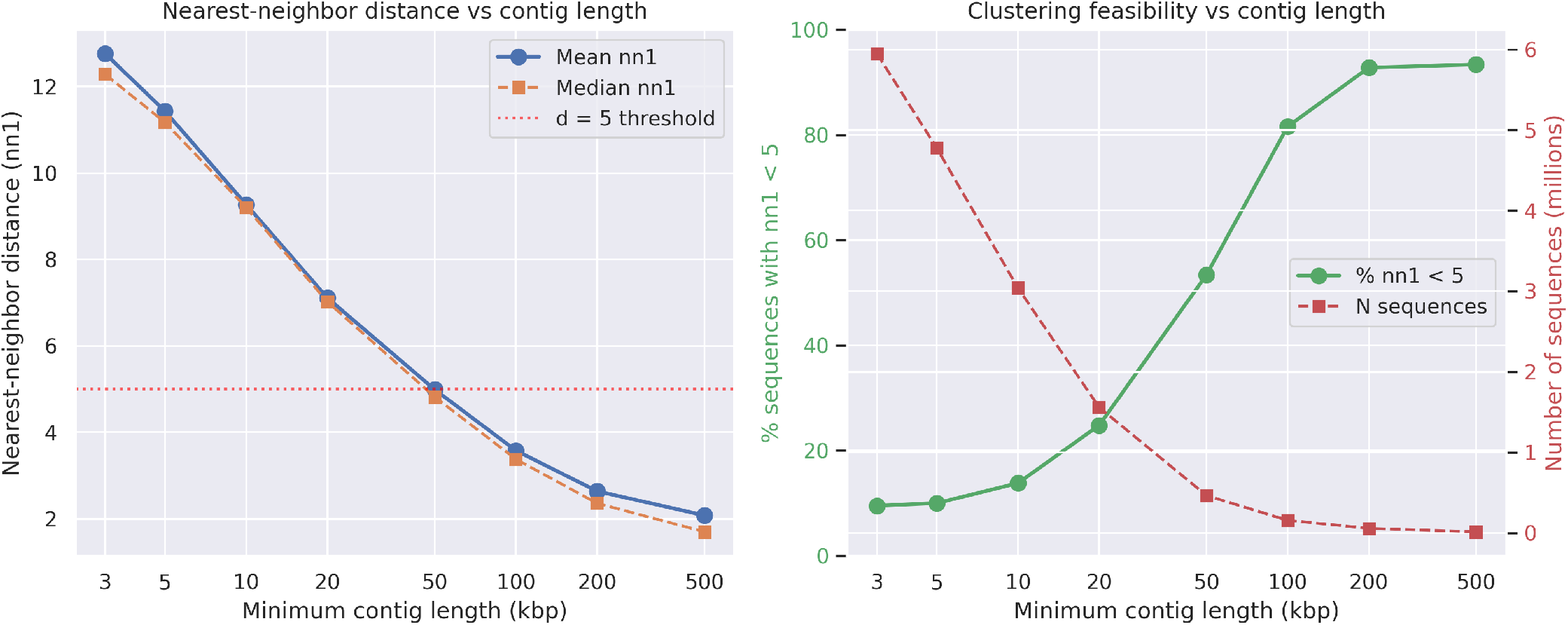
Nearest-neighbor distance (left) and clustering feasibility (right) vs minimum contig length. Clear knee at 50-100 kbp.

#### Theoretical basis for the length threshold

The observed knee at 50–100 kbp has a theoretical basis in the statistics of k-mer sampling. Under a multinomial model, the coefficient of variation (CV) of a k-mer frequency estimate scales as 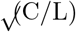, where C is the number of canonical k-mers at that k and L is the sequence length^7,8^. At a given sequence length, longer k-mers—which have more canonical types sharing the same total sequence—are estimated less precisely (Table 13):

**Table 13:** Predicted coefficient of variation (%) for k-mer frequency estimation at representative sequence lengths, assuming uniform k-mer distribution (multinomial sampling).

| k | Canonical types (C) | CV at 10 kbp | CV at 50 kbp | CV at 100 kbp | CV at 500 kbp |
| --- | --- | --- | --- | --- | --- |
| 1 | 2 | 1.4% | 0.6% | 0.4% | 0.2% |
| 2 | 10 | 3.2% | 1.4% | 1.0% | 0.4% |
| 3 | 32 | 5.7% | 2.5% | 1.8% | 0.8% |
| 4 | 136 | 11.7% | 5.2% | 3.7% | 1.6% |
| 5 | 512 | 22.6% | 10.1% | 7.2% | 3.2% |
| 6 | 2,080 | 45.6% | 20.4% | 14.4% | 6.4% |

At 100 kbp, the dominant 6-mer features (2,080 canonical types, comprising 75% of the input vector) have a predicted CV of ~14%—each canonical 6-mer is observed approximately 48 times on average. Lower-order k-mers are estimated far more precisely: 5-mers at ~7% CV, 4-mers at ~4%, and 1–3-mers at <2%. A naive threshold for all 6-mers to reach CV < 10% would require ~208 kbp, yet the empirical clustering knee falls at 50–100 kbp. Several factors explain why the embedding tolerates higher hexamer noise than this naive prediction suggests. First, the multi-scale architecture provides complementary information: even when individual 6-mer estimates are noisy, the 692 lower-order features (k = 1–5) are precisely estimated and collectively constrain the latent representation. Second, the CLR transformation converts frequencies to log-ratios, which are more robust to proportional noise than raw frequencies. Third, the VAE’s bottleneck (2,772 → 384 dimensions) acts as a denoiser, projecting noisy high-dimensional observations onto a smooth low-dimensional manifold that preserves systematic compositional differences while averaging out sampling fluctuations. Fourth, natural k-mer distributions are far from uniform—some hexamers are an order of magnitude more frequent than others—so the information-carrying k-mers are estimated more precisely than the uniform-distribution prediction implies.

The CV predictions also explain why 10 kbp contigs are noise-dominated: at this length, 6-mer CV reaches 46% (each canonical 6-mer observed fewer than 5 times on average), 5-mer CV is 23%, and even tetranucleotides—the standard features for metagenomic clustering tools—have a CV of 12%. The transition from noise-dominated to signal-dominated regimes between 10 and 100 kbp thus reflects the progressive stabilization of k-mer frequency estimates, with the multi-scale architecture enabling effective clustering at shorter lengths than any single k-mer size would permit.

#### MCL clustering quality across length thresholds

At 100 kbp with d = 5, MCL produced consistently tight clusters (Table 14): the three largest communities had GC spans of just 4 percentage points, indicating genuine taxonomic coherence. Even at the more permissive inflation I = 1.4 (at d = 7, SFE_SE_5 model), GC spans widened to 12–18 pp—substantially looser, but still far tighter than Leiden’s 33–49 pp on a 10 kbp graph at d = 10, where MCL achieved 7–9 pp on the same graph. However, 50 kbp data showed inconsistent quality (some communities reaching 9–10 pp), confirming that 50 kbp is a marginal threshold where the noise-to-signal ratio prevents reliable community detection.

**Table 14:** GC content spans (pp) of the three largest MCL clusters at each length threshold and configuration.

| Threshold | Model | Config | C1 | C2 | C3 | Verdict |
| --- | --- | --- | --- | --- | --- | --- |
| 10 kbp | SFE_SE_5 | d = 5, I = 2.0 | 5 | 5 | 5 | Good quality, 12% coverage |
| 10 kbp | SFE_SE_5 | d = 7, I = 3.0 | 6 | 4 | 7 | Good quality, 24% coverage |
| 10 kbp | SFE_SE_5 | d = 10, I = 3.0 | 9 | 7 | 7 | Acceptable, 47% coverage |
| 50 kbp | SFE_SE_5 | d = 7, I = 3.0 | 6 | 8 | 5 | Marginal (wider variation beyond top 3) |
| **100 kbp** | **NCBI_5** | **d = 5, I = 3.0** | **4** | **4** | **4** | **Consistently tight** |
| 100 kbp | NCBI_5 | d = 5, I = 5.0 | 4 | 7 | 5 | Tight, smaller clusters |

#### NCBI_5 matches domain-specific training for clustering

Having identified 100 kbp as the most effective length threshold, we compared the two leading models— SFE_SE_5 (domain-specific brackish training) and NCBI_5 (reference genome training)—on the same downstream clustering task. Both used identical graph construction (in-degree capped at 100, d < 5, weights 1/(d + 0.1)) and MCL parameters (Table 15).

**Table 15:**
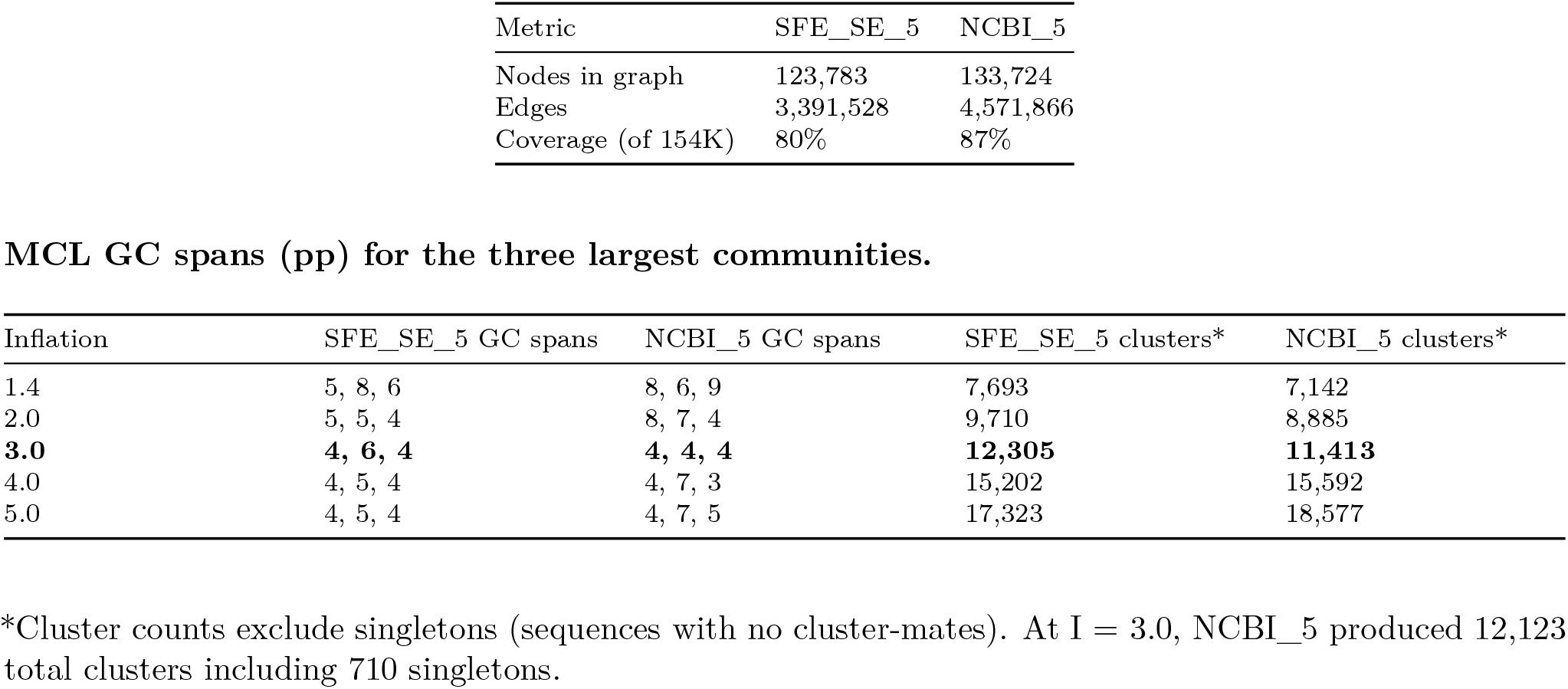
Clustering comparison of NCBI_5 vs SFE_SE_5 at 100 kbp.

| Metric | SFE_SE_5 | NCBI_5 |
| --- | --- | --- |
| Nodes in graph | 123,783 | 133,724 |
| Edges | 3,391,528 | 4,571,866 |
| Coverage (of 154K) | 80% | 87% |

MCL GC spans (pp) for the three largest communities.
| Inflation | SFE_SE_5 GC spans | NCBI_5 GC spans | SFE_SE_5 clusters* | NCBI_5 clusters* |
| --- | --- | --- | --- | --- |
| 1.4 | 5, 8, 6 | 8, 6, 9 | 7,693 | 7,142 |
| 2.0 | 5, 5, 4 | 8, 7, 4 | 9,710 | 8,885 |
| **3.0** | **4, 6, 4** | **4, 4, 4** | **12,305** | **11,413** |
| 4.0 | 4, 5, 4 | 4, 7, 3 | 15,202 | 15,592 |
| 5.0 | 4, 5, 4 | 4, 7, 5 | 17,323 | 18,577 |

\*Cluster counts exclude singletons (sequences with no cluster-mates). At $I = 3.0$ , NCBI\_5 produced 12,123 total clusters including 710 singletons.

At I = 3.0, NCBI_5 produced GC spans of 4 pp for all three largest clusters, while SFE_SE_5 produced 4, 6, and 4 pp. NCBI_5 connected nearly 10,000 more sequences (87% vs 80% coverage), producing a denser graph with 35% more edges. This is notable given that NCBI_5 was trained exclusively on prokaryotic reference genomes, while SFE_SE_5 was trained on 4.8 million domain-specific brackish contigs.

Despite NCBI_5’s substantial Spearman advantage over SFE_SE_5 on both full brackish data (+0.145; 0.837 vs 0.692 mean) and 100 kbp brackish data (+0.070; 0.836 vs 0.766), both models achieved comparable MCL cluster quality—reinforcing the dissociation between proxy metrics and downstream performance noted above (Table 9).

#### The 100 kbp configuration

At 100 kbp with distance threshold d = 5 and MCL inflation I = 3.0, using the NCBI_5 model we obtained our best clustering result:

- **154**,**041 contigs** (SFE+SE, ≥ 100 kbp)
- **87% connected** (133,724 nodes) in the in-degree-capped k-NN graph at d = 5
- **12**,**123 MCL clusters** (11,413 with two or more members; Figure 4B), largest containing 182 sequences
- **GC spans of 4 pp** for the three largest clusters; across all 11,413 non-singleton clusters, the median GC span was 1.2 pp and 95% of clusters had spans ≤ 5 pp (maximum 12.9 pp)
- **Intrinsic dimensionality** d-hat = 3.74 (clean, low-dimensional manifold)
- All three graph construction variants (symmetric, mutual, capped) produced broadly similar structure at this threshold

The distance threshold d = 5 matches the natural neighborhood structure of the 100 kbp data (mean nn1 = 3.58, 81.6% of sequences have a neighbor within d = 5). At this threshold, even Leiden achieved considerably tighter clusters than on the 10 kbp graph:

#### Leiden vs MCL GC spans (pp) at 100 kbp d = 5, in-degree capped graph (NCBI_5 model)

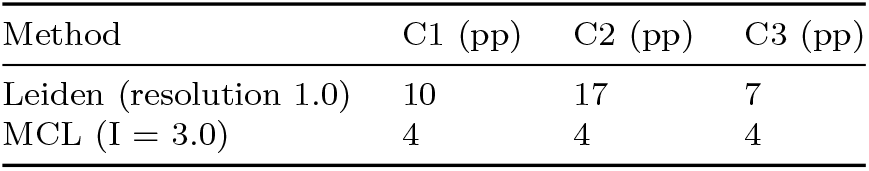

Leiden’s GC spans of 7–17 pp at 100 kbp represent a substantial improvement over its 33–49 pp on the 10 kbp graph, reflecting fewer spurious hub-mediated edges at this threshold. Nevertheless, MCL still produced tighter clusters (GC spans up to ~4× narrower). Leiden’s performance was insensitive to the resolution parameter: across a 10-fold range (0.2–2.0), it produced nearly identical partitions (5,779–5,872 clusters, <2% variation; see Methods).

### Taxonomic Validation Confirms Cluster Coherence

The tight GC content spans presented above provide indirect evidence of cluster quality, but direct taxonomic validation is essential. We performed a multi-phase annotation of the 133,724 contigs in the MCL graph (I = 3.0, d = 5, 100 kbp), combining five independent classification methods plus cluster-based propagation to achieve 96.8% annotation coverage.

#### NCBI signpost analysis

We embedded 175,213 NCBI RefSeq representative genome contigs (≥ 100 kbp) through the frozen NCBI_5 encoder and identified brackish contigs within Euclidean distance d < 5 of any reference sequence. Only 7,348 NCBI contigs (4.2%) fell within this threshold—consistent with the expectation that most environmental organisms lack close reference genomes—but these mapped to 325 of 12,123 MCL clusters (2.7%), transferring taxonomy to 11,479 brackish contigs (8.6% of the graph). At the phylum level, consensus within clusters was 99.9%, and 80% of assigned contigs received taxonomy to genus or species depth. The high within-cluster agreement (99.9% at phylum) indicates that proximity-based taxonomy transfer from reference genomes to environmental contigs produces coherent assignments.

#### Independent validation by GTDB-Tk

To validate these assignments using a fundamentally different approach, we ran GTDB-Tk v2.6.1^9^ (marker gene phylogenetics against GTDB r226^10^) on all 154,041 contigs. Of these, 26,841 (17.4%) were classified with sufficient marker genes for phylogenetic placement (26,162 of which fell within the MCL graph); 80,893 (52.5%) contained markers but could not be placed in the reference tree; and 46,307 (30.1%) contained no detectable markers. For the 2,868 contigs with assignments from both the NCBI signpost method and GTDB-Tk, agreement was 100% at domain level and 99.9% at both phylum and class levels after mapping between NCBI and GTDB nomenclatures—only 2 genuine phylum-level and 3 class-level disagreements across the entire dataset.

#### Protein-level validation by MMseqs2

As a third independent validation, we classified all contigs using MMseqs2 taxonomy^11,12^ against the NCBI nr database (see Methods). Of 154,041 contigs, 102,771 (66.7%) received phylum-level consensus (95,563 of which fell within the MCL graph)—3.8-fold more than GTDB-Tk (17.4%)—reflecting the broader coverage of protein homology search across all predicted ORFs rather than a restricted set of marker genes.

For the 11,050 contigs with both NCBI signpost and MMseqs2 phylum assignments, agreement was 99.7%— validating the signpost approach through a completely independent method based on protein homology rather than k-mer composition. Cluster purity from MMseqs2 assignments alone was 99.1% at phylum level across clusters with ≥ 2 classified members, closely matching the 99.2% from GTDB-Tk.

MMseqs2 also provides sub-domain taxonomy for eukaryotes—a capability absent from both GTDB-Tk (prokaryotes only) and Tiara (domain-level only): 15,091 contigs were classified as Eukaryota with phylum-level resolution, revealing lineage assignments for much of the eukaryotic content that other methods could only identify at domain level.

#### Cluster purity

Using GTDB-Tk classifications, we assessed taxonomic purity across MCL clusters containing two or more classified members (denominators vary by rank because not all clusters have classifications at finer taxonomic levels):

Purity remained above 98% through family level, declining only modestly to 95.6% at genus (Figure S1). Only 21 clusters exhibited phylum-level impurity. These purity estimates apply to the 2,753 clusters (23% of 12,123 total) that contain two or more GTDB-Tk-classified members—a subset restricted to prokaryotic contigs with sufficient marker genes for phylogenetic placement. The remaining clusters, which include eukaryotic, viral, and novel prokaryotic content, could not be assessed by this metric. Nevertheless, the near-perfect purity across all assessable clusters indicates that the MCL clusters correspond to genuine taxonomic units with high coherence.

To assess whether the high purity reflects over-splitting, we computed the complementary metric: for each taxon with ≥ 2 GTDB-Tk-classified contigs in the graph, the fraction residing in the largest cluster (Table 16b).

**Table 16:**
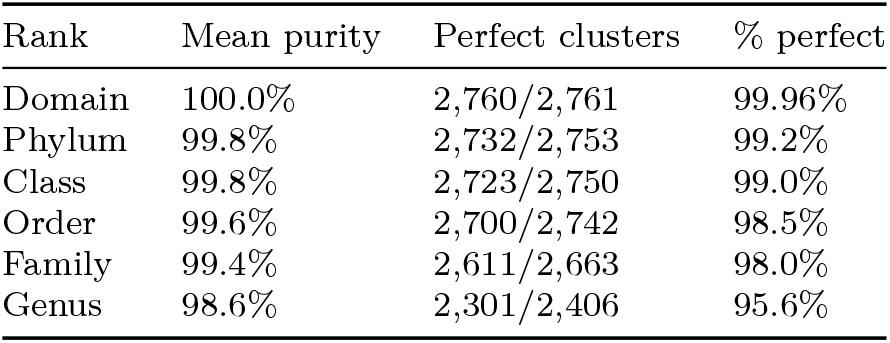
Taxonomic purity of MCL clusters by rank (GTDB-Tk classifications).

| Rank | Mean purity | Perfect clusters | % perfect |
| --- | --- | --- | --- |
| Domain | 100.0% | 2,760/2,761 | 99.96% |
| Phylum | 99.8% | 2,732/2,753 | 99.2% |
| Class | 99.8% | 2,723/2,750 | 99.0% |
| Order | 99.6% | 2,700/2,742 | 98.5% |
| Family | 99.4% | 2,611/2,663 | 98.0% |
| Genus | 98.6% | 2,301/2,406 | 95.6% |

**Table 16b:** Taxonomic completeness by rank (GTDB-Tk classifications).

| Rank | Mean completeness | Perfect taxa | % perfect |
| --- | --- | --- | --- |
| Species | 91.7% | 1,056/1,359 | 77.7% |
| Genus | 80.7% | 556/1,009 | 55.1% |
| Family | 69.7% | 219/574 | 38.2% |

Species-level completeness averaged 91.7%, with 77.7% of species contained entirely within a single cluster. Completeness naturally decreases at higher ranks, where each taxon spans many biologically distinct clusters. The most fragmented species were members of *Polynucleobacter* and *Pelagibacter*—genera known for extensive microdiversity—where splitting across 5–7 clusters likely reflects real strain-level compositional variation across the 32 metagenomes spanning two geographically distant environments. Both purity and completeness are computed on the same GTDB-Tk-classified subset and share its restriction to prokaryotic contigs with sufficient marker genes for phylogenetic placement.

#### Combined annotation coverage

Leveraging the high cluster purity, we propagated GTDB-Tk classifications to unclassified cluster members using majority vote with an 80% agreement threshold and minimum two classified members per cluster. We also applied geNomad^13^ for virus and plasmid detection and Tiara^14^ for eukaryotic domain classification, yielding a comprehensive annotation (Table 17; Figure S3):

**Table 17:** Combined annotation coverage of the MCL graph.

| Method | Contigs | % of graph |
| --- | --- | --- |
| NCBI signposts | 11,479 | 8.6% |
| GTDB-Tk (direct) | 26,162 | 19.6% |
| GTDB-Tk (propagated) | 56,794 | 42.5% |
| MMseqs2 (phylum consensus) | 95,563 | 71.5% |
| geNomad (virus/plasmid) | 18,628 | 13.9% |
| Tiara (eukaryote/organelle) | 18,832 | 14.1% |
| **Any annotation** | **129,483** | **96.8%** |

GTDB-Tk propagation was justified by the 99.2% phylum purity demonstrated above—assessed using only direct GTDB-Tk classifications, not propagated labels—and geNomad had zero overlap with GTDB-Tk (viral contigs lack prokaryotic marker genes), making the methods fully complementary. MMseqs2 resolved 76% of this gap: of the 17,540 contigs previously unannotated by the other five methods, 13,299 received MMseqs2 phylum-level classification—predominantly prokaryotic contigs that GTDB-Tk could not place due to insufficient marker gene alignment. Only 4,241 contigs (3.2%) remained unannotated—representing the most divergent or novel sequences in the dataset.

### The Embedding Captures Eukaryotic Biology That Standard Tools Cannot Classify

#### The eukaryotic dark matter

Among the annotation methods above, Tiara classification revealed that 25,247 contigs (16.4% of all 154,041 contigs ≥ 100 kbp) are eukaryotic. This population was almost entirely invisible to our other annotation tools: eukaryotic contigs constituted just 0.1% of contigs annotated by GTDB-Tk or NCBI signposts, but 18.5% of unannotated contigs—the single largest component of the “dark matter” (sequences escaping classification by standard tools) in the MCL graph. The explanation is straight-forward: GTDB-Tk uses prokaryotic marker gene sets, the NCBI signposts are predominantly prokaryotic, and geNomad detects viruses and plasmids. None of these methods is designed to identify eukaryotic content. This 16.4% contig fraction overstates eukaryotic species prevalence: at current assembly quality, larger eukaryotic genomes fragment into more contigs above the 100 kbp threshold than smaller prokaryotic genomes. This assembly fragmentation bias diminishes with improving long-read sequencing depth and read length; in the limit of complete assembly, the overrepresentation reduces to chromosome count differences.

#### Coherent eukaryotic clusters

Of the 18,365 eukaryotic contigs in the MCL graph, 17,829 (97.1%) fell into 2,746 eukaryote-dominated clusters, with the top 15 largest all showing 99–100% eukaryotic composition by Tiara classification. The remaining 536 eukaryotic contigs were distributed across prokaryote-dominated clusters. Only 17 clusters exhibited substantial eukaryotic–prokaryotic mixing, indicating clean separation between domains of life. This separation persisted despite NCBI_5’s training data consisting entirely of prokaryotic genomes.

#### Giant viruses and coding density validation

Cross-tabulation of Tiara and geNomad classifications identified 4,142 contigs in the MCL graph classified as both eukaryotic (by Tiara) and viral (by geNomad)— consistent with giant viruses (Nucleocytoviricota), which share eukaryotic compositional features while being detected as viral by marker gene methods^15^. This count represents individual contigs, not assembled genomes—a single giant virus genome may contribute multiple contigs ≥ 100 kbp—drawn from 32 metagenomes spanning 24 sampling sites across the San Francisco Estuary (8 sites, two seasons) and a transect across the Baltic Sea (16 sites). Coding density analysis (pyrodigal^16^, a Python implementation of Prodigal^17^) independently corroborated three populations with distinct gene organization: prokaryotes (median coding density 0.888), eukaryotes (0.748), and giant virus candidates (0.870), with giant viruses occupying an intermediate position consistent with their mosaic genomes. Because pyrodigal is a prokaryotic gene caller that cannot predict eukaryotic genes with introns, the eukaryotic value (0.748) reflects the fraction of sequence covered by prokaryotic-style open reading frames rather than true eukaryotic gene density; the lower value is expected precisely because intron-containing genes are fragmented by prokaryotic gene prediction.

## Discussion

The central finding of this work is that a general-purpose compositional embedding—trained once, without sample-specific retraining—can organize hundreds of thousands of metagenomic contigs into biologically coherent clusters spanning all domains of life. A single 384-dimensional latent space supports similarity search and graph-based clustering from the same representation, establishing that compositional information alone— captured through multi-scale k-mer frequencies—is sufficient to produce taxonomically coherent clusters (99.2% phylum purity) at the scale of modern long-read metagenomes.

Our approach offers several advantages over existing methods for metagenomic sequence analysis. By using multi-scale k-mer frequencies spanning 1-mers through 6-mers (2,772 features), the embedding captures compositional information across multiple scales—from nucleotide bias through codon usage to oligonucleotide signatures—providing 20-fold more features than the 136 tetranucleotide frequencies typically used for metagenomic sequence analysis. Requiring only sequence composition, the embedding applies to any individual contig without multi-sample experimental designs. A single set of 384-dimensional vectors supports nearest-neighbor retrieval and graph-based clustering via efficient HNSW indexing^1^, without retraining for each application. The finding that Euclidean distance outperforms cosine in this latent space (+0.076 Spearman) has a straightforward explanation: because the MSE loss penalizes L2 distance in the reconstruction space, Euclidean proximity in the latent space reliably reflects compositional similarity. This validates the use of standard L2-based vector databases and approximate nearest-neighbor algorithms without modification. This embedding approach is fundamentally distinct from metagenomic binning tools such as VAMB and SemiBin. The relevant comparison for a composition-only embedding is against other composition-only dimensionality reduction methods, which is why we benchmark against PCA and raw CLR baselines at matched dimensionality.

More fundamentally, the embedding captures global compositional similarity—something alignment-based methods do not capture. Genomic composition is approximately consistent across a genome^2^, so non-overlapping contigs from the same organism map to nearby latent points even when they share no local sequence homology; alignment-based clustering, by contrast, requires shared homologous regions and would treat such contigs as unrelated. Because the embedding places all contigs in a shared space regardless of sample origin, it naturally supports cross-environment comparison: in this study, contigs from 32 metagenomes spanning two geographically distant brackish environments co-clustered by taxonomic identity rather than by sample. This cross-sample organization—grouping related sequences regardless of origin—is precisely the capability that compositional clustering provides: it accommodates the genomic variation within natural populations rather than forcing it into a consensus, enabling higher-level ecological comparison that per-sample genome reconstruction cannot.

Several methodological findings from this work have implications for the broader field of learned representations in genomics. Our results challenge the assumption that reconstruction loss—a commonly reported performance metric for VAE-based metagenomic methods—reliably indicates embedding quality. The model with the lowest reconstruction MSE produced the worst embedding by Spearman correlation, while a model with 1.3× higher MSE produced a better embedding (Table 9). Spearman correlation serves as an effective screening metric—it correctly identifies that focused training substantially outperforms mixed training (0.812 vs 0.644 on common test data; Table 9)—but it does not predict clustering quality among comparably good embeddings: NCBI_5 and SFE_SE_5 produced comparable MCL cluster quality despite NCBI_5’s substantial Spearman advantage on both full brackish data (+0.145; 0.837 vs 0.692) and 100 kbp brackish data (+0.070; 0.836 vs 0.766; Table 8), and the practical difference between the two models lay in graph coverage (87% vs 80%)—a property neither metric captured. This dissociation underscores that the relevant question is not whether the embedding perfectly preserves k-mer distances, but whether it produces clusters that correspond to biological groups. This dissociation extends beyond model comparisons: PCA-384, a linear baseline, achieves higher Spearman correlation (0.948 vs 0.837) and tighter GC content spans (median 0.98 vs 1.16 pp), yet produces taxonomically less coherent clusters—with 11.5 percentage points fewer single-cluster species (66.4% vs 77.9%) and lower ARI, V-measure, and pairwise F1 at genus and species level (Tables 10–11). The VAE’s learned nonlinear representation trades distance fidelity for clusterable structure, a trade-off that linear dimensionality reduction cannot make.

Training data properties mattered more than scale. A curated dataset of 656,000 prokaryotic reference genome contigs with longer length distribution (median ~37 kbp vs ~8 kbp), complete genomes rather than fragmentary environmental contigs, and broad taxonomic coverage (~20,000 representative genomes) outperformed 6.7 million domain-specific brackish sequences on embedding quality and matched them for clustering quality—a 10-fold reduction in training data. Because these properties are confounded, we cannot attribute the advantage to any single factor; rather, the combination of curation, length distribution, and compositional coherence appears to outweigh sheer scale. Indeed, naively combining the brackish and reference genome training sets actively degraded performance (Spearman 0.649 for the combined model vs 0.837 and 0.766 for NCBI_5 and SFE_SE_1 separately), suggesting that mixing data sources with different distributional properties can create conflicting optimization targets.

A possible explanation for the effectiveness of small, curated datasets lies in the VAE’s training dynamics (see Methods). Because the VAE’s regularization and stochastic sampling together cause each training sequence to define a neighborhood rather than a point in latent space, the decoder must generalize across perturbations rather than memorize fixed mappings. Over 1,000 epochs, each of NCBI_5’s 656,000 sequences is perturbed a thousand times, allowing even a relatively small number of clean, complete-genome training points to populate the latent space with smooth, coherent regions. Noisier or more heterogeneous training data may produce less consistent neighborhoods, which may explain why more data does not necessarily yield a better embedding.

The choice of community detection algorithm proved critical. Modularity-based methods such as Leiden^3^ are widely used in single-cell and network biology, but our results show they are poorly suited to the nearest-neighbor graphs that arise from metagenomic sequence search, where hub nodes often form. At 100 kbp, MCL produced clusters with GC content spans of 4 percentage points (pp) compared to 7–17 pp for Leiden. The hub problem—where highly connected sequences bridge compositionally unrelated clusters—is inherent to nearest-neighbor graphs in high-dimensional spaces, where geometric concentration^4^ causes certain points to appear as nearest neighbors of disproportionately many others. MCL’s flow-based approach^5,6^ is naturally robust to this topology because its iterative inflation step progressively attenuates weak indirect connections, a property that modularity optimization lacks. These results suggest that MCL is well suited to nearest-neighbor graphs constructed from metagenomic embeddings, where hub nodes commonly arise.

Beyond the choice of clustering algorithm, the embedding itself reveals biological structure consistent with current understanding of microbial community ecology. GC content emerges as the primary axis of variation, reflecting the fundamental role of base composition in shaping k-mer frequencies through mutation bias and selection^7^. Beyond GC content, the latent space captures additional compositional axes: at 100 kbp, the intrinsic dimensionality of approximately four^8^ suggests that species-level discrimination relies on GC content plus a small number of orthogonal compositional features, likely reflecting dinucleotide biases and codon usage preferences. The archipelago structure—discrete islands separated by empty space rather than a continuum of gradual transitions—reflects the compositional discreteness of microbial species: each lineage occupies a narrow region of k-mer space, and the embedding preserves these gaps rather than interpolating between them. Lineages sampled deeply enough to produce multiple assembled contigs form dense islands, while those contributing too few contigs appear as isolated singletons.

Sequence length is the primary driver of clustering quality, with a clear knee at 50–100 kbp: below this threshold, the majority of sequences are isolated singletons, while above it, the data forms a clean low-dimensional manifold amenable to graph-based community detection. This length dependence has a quantitative basis in multinomial sampling statistics^9,10^: the coefficient of variation of k-mer frequency estimates scales as 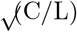, predicting ~14% CV for hexamers at 100 kbp but ~46% at 10 kbp (Table 13). Although a naive threshold of ~208 kbp would be required for all canonical hexamers to reach CV < 10%, the multi-scale architecture, CLR transformation, and VAE denoising collectively lower the practical threshold to 50–100 kbp. The 100 kbp threshold that enables effective clustering is practical only because long-read sequencing technologies routinely produce contigs of this length. Long-read assemblies yield substantially more contigs above this threshold than short-read assemblies; in short-read assemblies, most contigs fall in the noise-dominated regime. As long-read sequencing continues to improve—producing longer contigs—a growing fraction of the metagenome will become accessible to compositional clustering.

Taxonomic validation confirmed the biological coherence of these clusters. Independent marker gene phylogenetics (GTDB-Tk^11^) and protein-level homology search (MMseqs2^12^) both yielded >99% phylum-level cluster purity among the clusters assessable by each respective method (Table 16; MMseqs2 purity reported in text), despite the embedding containing no phylogenetic information—indicating that compositional proximity in the latent space corresponds to phylogenetic relatedness. The complementary metric, completeness, confirms that high purity does not merely reflect over-splitting: species-level completeness averaged 91.7%, with 77.7% of species contained entirely within a single cluster (Table 16b). The most fragmented species belonged to *Polynucleobacter* and *Pelagibacter*—genera known for extensive microdiversity—where splitting likely reflects real compositional variation across environments. The largest component of previously uncharacterized sequences proved to be eukaryotic: Tiara^13^ domain classification identified one in six contigs as eukaryotic by contig count (assembly fragmentation inflates this fraction relative to species prevalence). Cross-referencing with geNomad^14^ virus detection revealed a distinct population of giant virus candidates (Nucleocytoviricota^15^) with intermediate coding density between eukaryotes and prokaryotes. These findings demonstrate that the VAE captures organizational principles extending well beyond its prokaryotic training data.

Several limitations of this work should be acknowledged. The model was trained exclusively on prokaryotic reference genomes, yet 16.4% of contigs in the target dataset are eukaryotic; while these sequences form coherent clusters, their organization has not been evaluated against eukaryotic ground truth. Effective clustering is limited to contigs of at least 100 kbp, which represents only a fraction of typical metagenome assemblies—the majority of contigs in our dataset fall below this threshold and remain unclustered. Our training and evaluation focused on brackish and reference genome sequences; whether these findings about training data properties generalize to other environments (soil, host-associated, extreme) remains untested, though the strong cross-domain performance of the NCBI-trained model is encouraging. Tiara’s domain classification, which we use to identify eukaryotic sequences, is itself based on k-mer frequencies, introducing some methodological circularity; however, the coding density analysis (pyrodigal, a prokaryotic gene caller) provides independent corroboration based on gene organization rather than k-mer composition, confirming the three-population structure. We evaluated embedding quality on held-out validation sets (10% of each training file) that also served as the monitoring set for learning rate scheduling, rather than on fully independent test data; however, the consistently small train/validation loss gaps (Tables 2–3) suggest minimal overfitting risk. The NCBI reference genomes used for training span a wide range of assemblers and sequencing platforms, and the model’s strong performance on Flye-assembled metagenomic data suggests robustness to assembler choice, but systematic evaluation across metagenomic assemblers has not been performed. Finally, the Spearman correlation values were computed from a single evaluation (100 queries × 50 neighbors); bootstrap confidence intervals over query resampling (10,000 resamples) confirmed stable estimates (e.g., NCBI_5 on brackish data: *ρ* = 0.831 [0.757, 0.880]), but the 100-query unit limits precision to approximately $±$0.06.

Several directions for future work emerge from these results. The absence of eukaryotic training data is the most obvious gap: eukaryotic contigs constitute the single largest unresolved component. Training on eukaryotic reference genomes, of which NCBI now catalogs over 21,000 species, could improve the organization of this fraction and better resolve the boundary between eukaryotic sequences and giant viruses that currently occupy overlapping regions of the latent space. Evaluation on metagenomes from other environments—soil, host-associated, hydrothermal—would establish the generality of these findings. Extending the cross-environment comparison to larger collections—hundreds of metagenomes spanning ecological gradients—could provide a sample-independent coordinate system for tracking community composition at scale.

By providing a general-purpose embedding that preserves local compositional similarity, this work enables scalable metagenomic analysis across studies. The same embedding supports similarity search, graph-based clustering, and comparative analysis—capabilities increasingly needed as environmental sequencing projects generate millions of contigs across hundreds of samples. The finding that small, well-curated training sets can match or outperform far larger datasets may have implications beyond metagenomics for embedding tasks where local distance structure matters. The archipelago structure of the latent space demonstrates that the compositional discreteness of microbial species is preserved through the embedding, while the taxonomic validation confirms that compositional embeddings capture biologically meaningful structure across all domains of life. Together, these results establish a foundation for organizing and querying metagenomic sequence space at scale.

## Code and Data Availability

The VAE training code, inference scripts, k-mer frequency calculator, and figure generation scripts are available at https://github.com/tnn111/Embedding. Trained model weights will be deposited in a public repository prior to journal submission. Raw sequencing data for the San Francisco Estuary metagenomes are available under BioProject accession [TBD]; Baltic Sea metagenomes under [TBD]. Microflora Danica assemblies are available from the European Nucleotide Archive as described in Sereika *et al*.^20^ NCBI RefSeq representative genomes were downloaded December 7, 2025.

---

**Figure S1:**
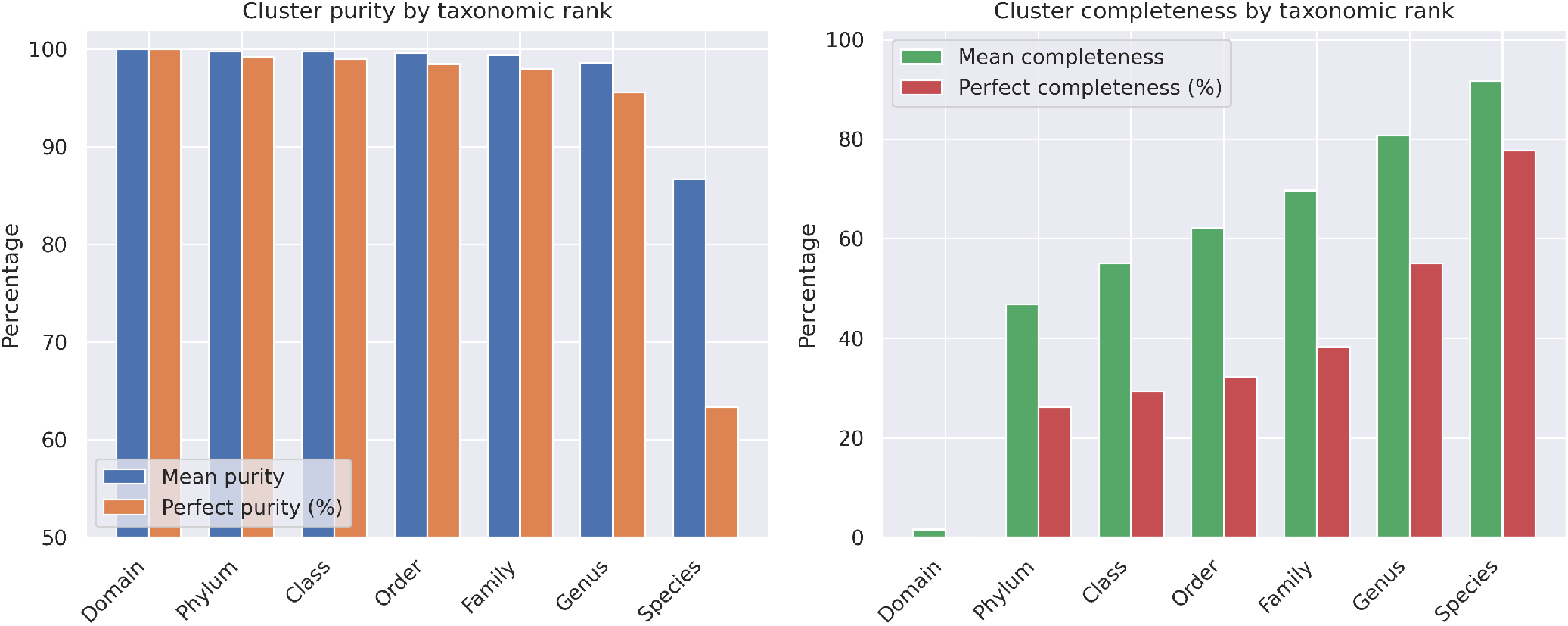
Cluster purity (left) and completeness (right) by taxonomic rank. GTDB-Tk validated.

**Figure S2:**
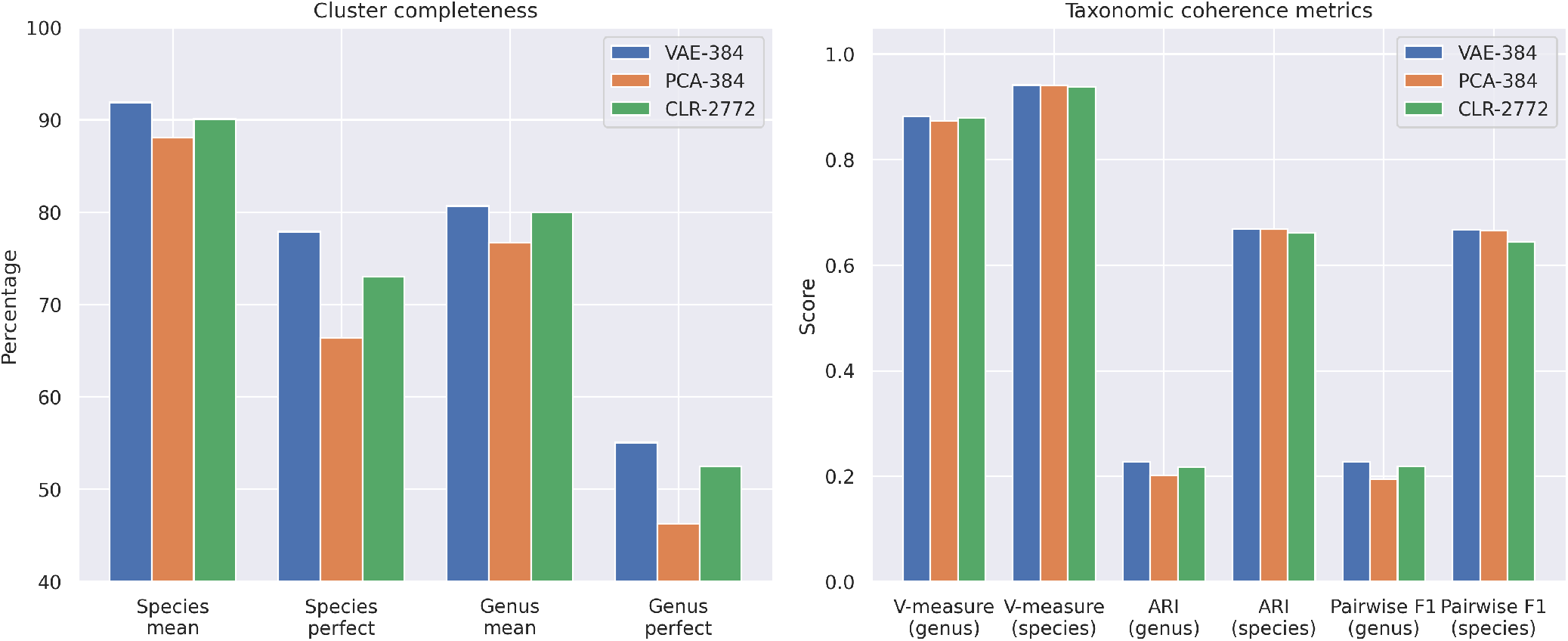
VAE vs PCA vs CLR baselines. Cluster completeness (left) and taxonomic coherence metrics (right).

**Figure S3:**
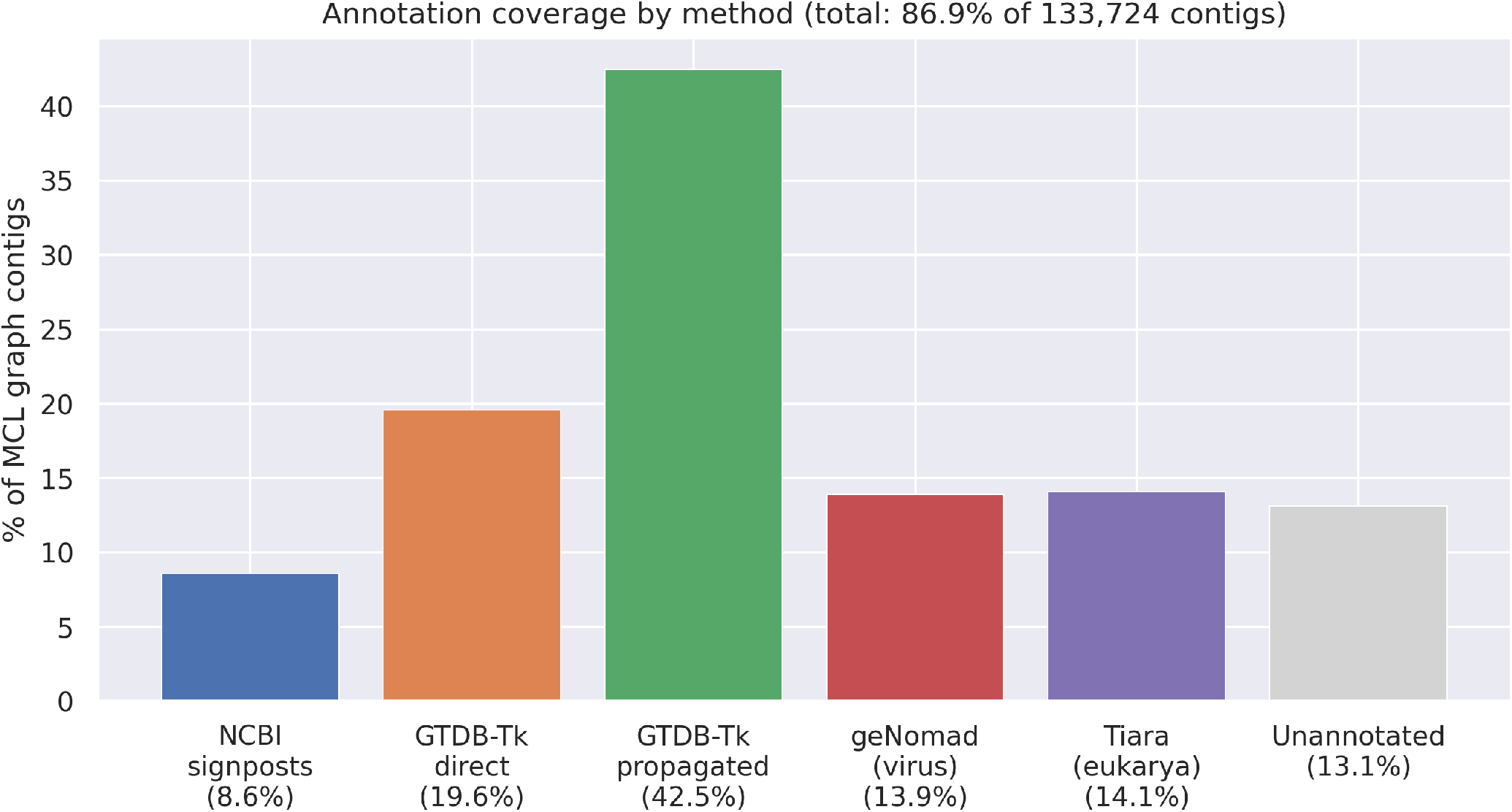
Annotation coverage by method. Five complementary methods achieve 86.9% coverage of the MCL graph.

